# nail: software for high-speed sequence annotation with profile hidden Markov models

**DOI:** 10.1101/2024.01.27.577580

**Authors:** Jack W. Roddy, David H. Rich, Travis J. Wheeler

## Abstract

*Fast is fine, but accuracy is final*.

--Wyatt Earp

**Background:** Profile hidden Markov models (pHMMs) deliver state-of-the-art sensitivity for sequence annotation, but the Forward/Backward algorithm fills a dynamic programming matrix sized by the product of model and sequence lengths, making pHMM search slower than fast heuristic alignment tools like MMseqs2 by an order of magnitude or more.

**Results:** We introduce `nail`, which approximates Forward/Backward by computing only a sparse cloud of high-probability matrix cells, recovering accurate pHMM scores, E-values, and alignments at a fraction of the cost. `nail` annotates the ~2.4 billion protein MGnify metagenomic dataset with all of Pfam in 73.9 hours on a single 48 core machine, recovering nearly all of HMMER’s recall advantage over MMseqs2, with run time ~8.7x faster than HMMER3. Detailed analysis of HMMER-only multi-domain hits suggests that many of these matches missed by nail are the result of accumulating score across short, often fragmentary alignments to repetitive regions of the target, consistent with spurious hits rather than genuine homology. We also derive a closed-form approximation for single-sequence E-value calibration, eliminating a per-model simulation step. `nail` is released under the open BSD-3-clause license at https://github.com/TravisWheelerLab/nail.

## Introduction

### Profile hidden Markov models for high sensitivity

The dominant method for accurate annotation of biological sequences is sequence database search, in which an unlabeled sequence is classified by *aligning* it to sequences in an established database. Structure-based search has gained prominence, but the structure of an unannotated protein is typically unknown and structure prediction is both slow and usually dependent on a first-pass database search to discover homologous sequences. Meanwhile, protein language models are promising, but provide limited interpretability and carry substantial false labeling risk [28, 29].

Alignment-based annotation has historically been associated with the Smith-Waterman algorithm [36] and fast heuristics; BLAST [1], in particular, has long been an important part of the annotation ecosystem. Profile hidden Markov models (pHMMs [24, 8, 9]) have been shown to produce superior sequence search sensitivity.

Much of the sensitivity of pHMMs is due to their natural representation of profiles – when a collection of sequence family members is used to train the model, a pHMM captures the position-specific letter and gap frequencies inherent to the family. Profile representation of a family of sequences allows for improved search sensitivity relative to search using a collection of individual sequences [19, 11, 23], and these families also enable faster annotation time when sequences can be compared to a single family profile rather than numerous family instances. This pair of benefits has driven the development and use of databases of sequence families and accompanying pHMMs all across bioinformatics, e.g. [16, 18, 25, 21, 4, 26, 38].

Unlike other alignment methods that compute just a single highest-scoring alignment, pHMMs naturally enable computation of support for homology based on the sum of the probabilities of *all* alignments via the Forward/Backward (FB) algorithm [30, 24]. In addition to producing a measure of match significance, FB produces posterior probabilities that two residues are aligned, enabling greater alignment accuracy [20, 7] as well as improved mechanisms for addressing composition bias and determining alignment boundaries [10].

Computing FB is computationally expensive: to align a pHMM to a sequence, FB requires completion of a dynamic programming matrix with size determined by the product of the the lengths of the pHMM and target sequence, with each matrix cell requiring additional calculations to capture the sum of alignment probabilities (see [11] for discussion). HMMER3 introduced a pipeline in which most candidates are never subjected to expensive FB analysis, thanks to a series of earlier filter stages [11]. The result was a tool with speed comparable to BLAST, but with superior sensitivity. In common use cases, the first filter of HMMER3 (called MSV) consumes ~70% of HMMER’s run time, while FB consumes ~20% of time and is primarily responsible for large memory usage due to the quadratic-sized dynamic programming matrix required for recovering the alignment. FB dominates run time in cases of queries with high length or large numbers of true matches, and becomes the primary run time bottleneck in the event of improved speed for the earlier filter phases [3, 28].

### Algorithms for high speed

Heuristic alignment tools have grown dramatically faster over the past decade, with the most sensitive of them now approaching BLAST-level recall on protein homology search at run times often more than an order of magnitude faster. MMseqs2 [37] is the current leader along this axis, supporting both single-sequence and profile queries. Its speed comes from an aggressive filtering pipeline: optimized lookup tables identify high-scoring spaced length-k matches between query and target, profile-sequence pairs with two co-diagonal matches are extended into ungapped alignments, and bounded gapped alignment with a profile (position-specific scoring) model is then performed on only the top-scoring above-threshold pairs (default 300). The pipeline terminates with a gapped Viterbi-style alignment and does not compute Forward/Backward, leaving a sensitivity gap relative to pHMM search [23]. Another widely-used tool in this space, DIAMOND [6], lacks profile query support and shows slightly lower benchmark sensitivity than MMseqs2 (see Figures 3 and 4, also [23]).

### A pipeline for high-speed <u>and</u> sensitive alignment

Here, we describe a sequence search pipeline that first rapidly identifies candidate sequence matches, then employs a fast FB heuristic to improve alignment recall. Candidate sequence matches are found using the MMseqs2 software suite tuned with custom parameters. The fast FB heuristic limits search space in the FB dynamic programming (DP) matrix to a high-probability cloud around the alignment seed generated by MMseqs2, and results in calculations that closely approximate the results of the full FB algorithm. The sparse FB implementation (Figure 1), along with downstream analyses making use of the resulting sparse posterior probability matrix, are adapted from HMMER3. The algorithm is implemented from scratch in the Rust programming language, producing a modern and stable codebase with reduced runtime and memory requirements for highly-sensitive sequence annotation. The software, called `nail` (for <u>n</u>ail is an <u>a</u>lignment <u>i</u>nference too<u>l</u>), supports search using either protein sequence or pHMM queries, and is released under an open (BSD 3-clause) license; source code is available at https://github.com/TravisWheelerLab/nail and is hosted on the official Rust package registry at https://crates.io/crates/nail.

**Fig. 1:**
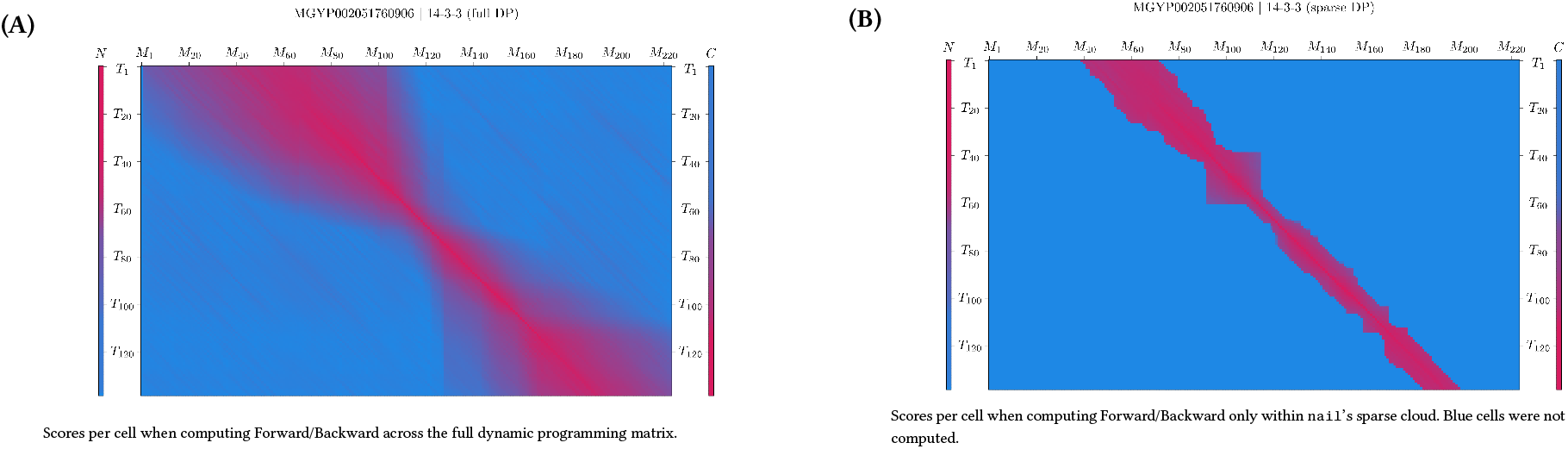
Sparsely filled Forward/Backward matrix captures most of the probability mass. Comparison of the Match state scores for full and sparse dynamic programming matrices for alignment of the MGNIFY sequence MGYP002051760906 versus the Pfam domain 14-4-3 (PF00244). Sequence positions are aligned along the y-axis. Model positions, along with terminal N and C states (left and right vertical bars, respectively) are aligned along the x-axis. Bright red colors correspond to high scores, and blue colors correspond to very low (or no) score. Score at a given cell is the sum of the Forward matrix value at that cell and the Backward matrix value at that cell; this corresponds to the posterior probability, prior to normalization (see Supplementary Materials for more). Though the heatmap shows high scores along a generally-diagonal path covering the full length of the target (from top to bottom), the maximum scoring alignment in fact skips the first 40 residues in the sequence (by emitting them from the N state) and the last 32 residue (by emitting them from the C state).

In the following sections, we demonstrate the efficacy of nail’s sparse FB implementation and explore the impact of the overall pipeline on speed and sensitivity on both a controlled benchmark and a large-scale annotation of the MGnify metagenomic database. Within the MGnify analysis, we characterize the structural patterns underlying HMMER’s apparent recall advantage on multi-domain hits, finding that a dominant category consists of many overlapping weak alignments to a small portion of the model, a signature more consistent with repetitive target sequence than with multiple genuine domain copies. We also present a closed-form approximation that replaces the simulation step in HMMER3’s single-sequence E-value calibration; this solution eliminates a substantial per-query computational cost and yields prediction error smaller than the seed-to-seed noise of the simulation itself. We then provide a thorough description of nail’s implementation.

## Results

The primary innovation of `nail` is the development of an approximate method that reduces the time and memory required for computation of the Forward/Backward (FB) algorithms for pHMMs, along with downstream analyses that are based on posterior probabilities resulting from FB (including creation of an alignment). The approach is a close cousin to the X-drop heuristic [41] used in BLAST: start with a seed that establishes a region of interest within the DP matrix, and perform DP calculations out in both directions until pruning conditions are met. Figure 1 presents a single example of the reduced computation required by `nail`’s sparse FB for a fairly short alignment of one Pfam-based pHMM against a sequence belonging to the family. In the current release of `nail`, seeds are acquired by running MMseqs2 as a subroutine. Details are provided in the Methods section.

We begin by describing the data and process used for evaluation, then demonstrate the space-pruning efficacy of `nail`’s Cloud Search approach. We then show that annotation with `nail` closes most of the accuracy gap between sensitive HMMER search and BLAST or MMseqs2 search, with only modest additional runtime over MMseqs2. Scripts and notes to reproduce benchmarking results can be found at https://github.com/TravisWheelerLab/nail-benchmarks.

### The nail pipeline – a sketch

The `nail` pipeline begins by identifying a set of candidate query/target sequence pairs by running the standard MMseqs2 search pipeline with a few parameters adjusted (see Table 3). The first and last positions of each surviving MMseqs2 alignment are used as seed cells in an implicit Forward/Backward alignment matrix. Using the seed cells as a starting point, a heuristic search algorithm (Cloud Search) identifies a contiguous subset of Forward/Backward matrix cells with non-negligible probability. Within this reduced set of matrix cells, `nail` then completes a sparse variant of Forward/Backward (FB), producing an overall alignment score along with position-specific posterior probabilities that positions are aligned; these posterior probabilities are used to compute a composition bias score adjustment along with the final sequence alignment. See Methods for more details.

### Benchmarking philosophy

In the analysis that follows, we rely on two summary statistics as the arbiter of sensitivity: *recall* and *coverage*. When a benchmark has ground truth (e.g. a set of sequences known to be homologous), we can discuss *recall*, which is the fraction of the known positives that are recovered. It is vital to consider recall in tandem with some measure of a false positive rate (FPR), which requires that the benchmark includes planted decoy sequences that are known to be not homologous to the query sequences. One strategy is to identify the E-value at which the number of false positives exceeds some threshold (e.g. 0.01) per search, and report the fraction of all true positive matches exceeding that score: the “*Recall* at 0.01 FP per search”. This summary statistic is easy to interpret and generally agrees with relative ordering of methods in analyses such as Figure 3, or those reported in papers introducing MMseqs2 [37] and DIAMOND [6]. Alternatively, recall may be viewed as a function of FPR, offering perspective on the sensitivity of a tool across a spectrum of tolerance for false annotation.

In the absence of ground truth (e.g. when annotating a new genome), we turn to *coverage*, which is the fraction of the target sequences receiving a score or E-value better than some *cutoff* value. In rough order of usefulness, *cutoffs* may be (i) manually-curated per query (e.g. Pfam domain “trusted cutoffs”), (ii) learned per query (e.g. ranking alignments to decoy sequences), or (iii) assigned as a blanket value (e.g. `score` ≥ 20 `bits` or `E-value` ≤ `1E-3`).

It is important to keep in mind that the benefits of sensitive search tools are evident among (almost exclusively) the most challenging search conditions. In other words, it is not interesting to find that two competing tools both report matches between highly similar sequences, but it is interesting when one tool recovers a distant homolog that the other tool fails to find. It follows that results comparing the sensitivity between search tools are of limited utility when the benchmarks are dominated by easy-to-find matches.

### A benchmark

The first few experiments were performed using a benchmark derived using multiple sequence alignments (MSAs) from Pfam v36.0 [26]

#### Definitions

𝔸: a source MSA

𝔸_Q_, 𝔸_T_: disjoint subsets of 𝔸

(q, t): a query–target sequence pair, (q, t) ∈ 𝔸_Q_ × 𝔸_T_

The construction of the benchmark is as follows:

- For each seed MSA 𝔸 in Pfam:
  - remove all sequences from 𝔸 that are shorter than length 150 (note: very short, very low-scoring sequences are *very* difficult to recover)
  - sample disjoint subsets 𝔸_Q_, 𝔸_T_ from 𝔸 such that:
    i. `pid` (q, t) ≤ 25% for all (q, t) ∈ 𝔸_Q_ × 𝔸_T_
    ii. 𝔸_T_ contains between 10 and 30 sequences (note: Pfam families are removed from consideration if it is not possible to create a 𝔸_T_ with at least 10 sequences; the cap of 30 sequences is intended to reduce the risk that a few families will dominate benchmarking results)
- The remaining target sequences from all Pfam domains comprise the set of trusted positives
- The target set is supplemented with shuffled sequences from Swissprot; these sequences are decoys. Decoys are conditionally sampled to match the length distribution of the true positives.

### Sparse Forward/Backward reduces computation, is a good approximation

To evaluate `nail`’s sparse FB method, we tested the extent to which it reduces the number of computed cells, as this directly impacts time and space utilization. This measurement largely relies on the Pfam-based benchmark described above. In order to highlight the extreme pruning efficacy for long sequences, the Pfam-derived set was supplemented with 6 pairs of very long protein sequences from Uniprot (Table 1). For each pair, one sequence was designated the *query* and the other the *target*.

**Table 1.**
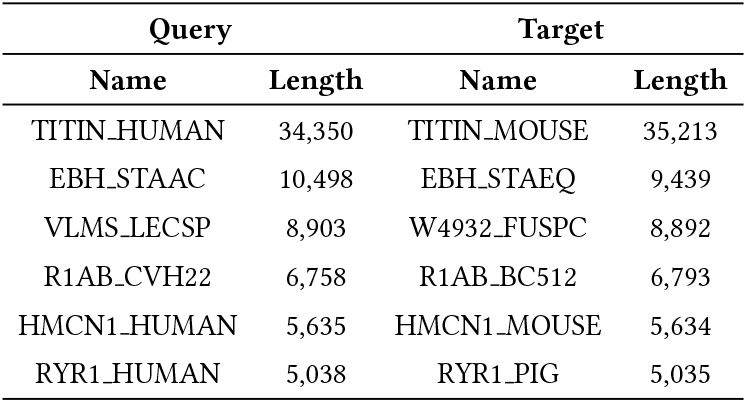
Long sequence pairs.

To analyze search space reduction, we calculated the percentage of the full quadratic search space that is computed by the sparse approach. In Figure 2A, matrix sparseness (y-axis, log-scale) is plotted against matrix size (x-axis, the product of the lengths of the query pHMM and target sequence; also log scale). Reduction in search space is modest for alignments of shorter sequences; this is not surprising, as the total size of the matrix is not particularly large, so that even a moderately-wide band around the maximum-scoring alignment will consume much of the full analysis space.

**Fig. 2:**
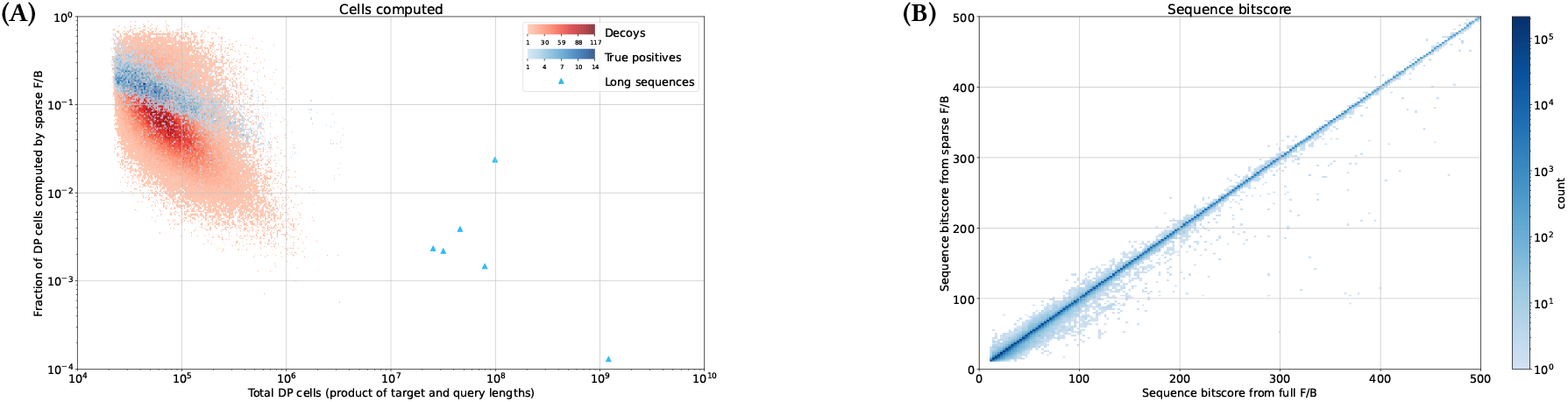
(A) Reduction in search space as function of size of that space. For each candidate alignment that survived the MMseqs2 Viterbi filter, we computed the fraction of the full DP matrix that was included in nail’s sparse cloud computations (y-axis). The darkness of each point in the plot indicates to the number of candidate alignments with the corresponding sparseness measure and number of cells in the full matrix (x-axis). Alignments of true domain matches are plotted in blue; red points show sparsification for false positive alignments; blue triangles (bottom right) show sparsification for long-sequence pairs, and follow the general trendline of space reduction as a function of matrix growth. **(B)** `nail` **’s sparse bit scores are accurate estimates of full bit scores**. Each point represents the relationship between sparse and full Forward scores for all (query, target) pairs identified in the MGnify/Pfam search. Loss of score shows up as vertical depression below diagonal. In some cases, a sparse alignment is reported with bias-adjusted score that is greater than the full matrix score; this is because `nail` implements a sparse variant of HMMER3’s heuristics for bias score adjustment, and matrix sparsity sometimes causes the bias-induced score adjustment to deviate.

**Fig. 3:**
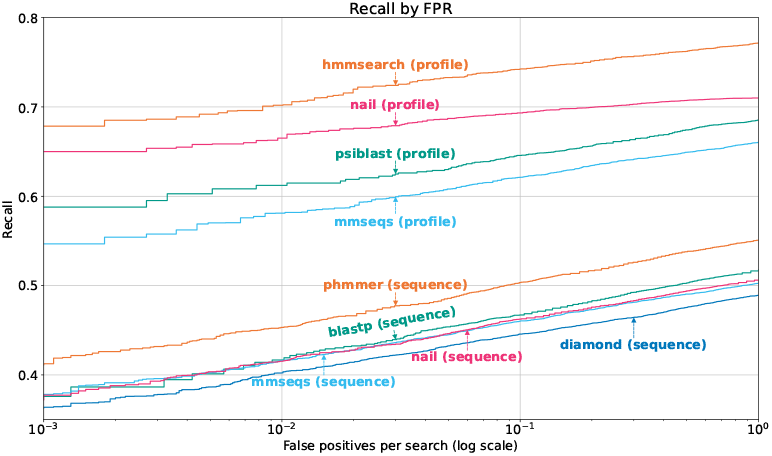
Recall as a function of false annotation rate. The benchmark consists of 11,187 true target sequences from 1,144 Pfam families, mixed with 1,118,700 shuffled sequences from Uniprot. `nail` variants were tested with `--mmseqs-s 12.0` and `--mmseqs-max-seqs 2000 --seed-mode` full; MMseqs2 variants were tested with `-s 12.0` and `--max-seqs 2000`; DIAMOND was tested with `--ultra-sensitive`; HMMER (hmmsearch, phmmer) and BLAST (psiblast, blastp) were tested with default parameters.

**Fig. 4:**
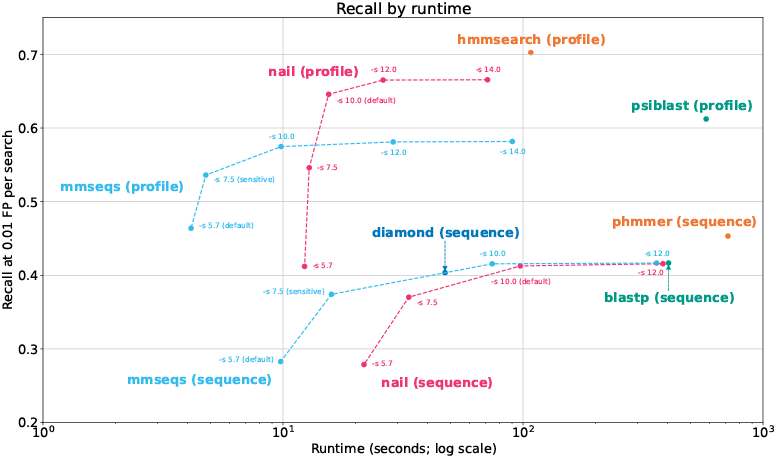
Runtime vs. recall. The Pfam-based benchmark was searched with tool variants to demonstrate performance-runtime tradeoffs. `nail` variants were tested with `--mmseqs-s` at various settings and `--mmseqs-max-seqs 2000 --seed-mode` full; MMseqs2 variants were tested with `-s` at various settings and `--max-seqs 2000`; DIAMOND was tested with `--ultra-sensitive`; HMMER (hmmsearch, phmmer) and BLAST (psiblast, blastp) were tested with default parameters. All tools were run with 48 threads.

For longer sequences (for example with a length-400 model aligned to a length-2500 protein, creating a matrix of size 10^6^), `nail`’s sparse method often restricts the total number of computed cells to 1% or less of the full size of the matrix; particularly large search space reductions are seen for alignment of the very long sequences described in Table 1 (blue triangles). Note that sparsification is greater on average for alignments involving decoy matches (Figure 2A, red dots), meaning that very little time is wasted computing the DP matrix for long non-homologous sequences.

We also measured how well the sparse analysis approximates alignment scores computed using full Forward/Backward. Figure 2B shows, for true positives from the domain benchmark, that the Forward score computed on the sparse matrix typically closely matches the score computed on the full matrix.

### Recall as a function of false annotation

We sought to determine how much of HMMER’s pHMM alignment accuracy is retained by `nail`’s sparse implementation, with MMseqs2 and blast included as a point of comparison. In the same evaluation, we explored the benefit of searching with profiles versus single sequence queries.

Each of the 1,144 query alignments was used to search for matching family members in the test database containing 11,187 real sequences mixed with 1,118,700 shuffled sequences. An alignment was considered to be ‘true positive’ if it matches the query from the same family. A hit that matched a shuffled (‘decoy’) sequence was defined as a ‘false positive’. Alignments between a query and target of differing families were treated as neutral (ignored) rather than being penalized, since it is possible that these matches correspond to previously unrecognized homology.

Figure 3 presents recall (fraction of all true positives that are recovered at a specific E-value cutoff) as a function of false annotation rate (number of false positive matches per query with that E-value or better). For each tested method, we pooled all alignments and sorted them by increasing E-value to produce a recall curve. The top collection of lines corresponds to search with a profile HMMER3’s hmmsearch tool was run with default settings (`-E` 10.0), establishing a sensitivity target. `nail` results are for default sensitivity settings ((‘-s 10 --maxseqs 1000’)). MMseqs2 was run with the same sensitive settings as default `nail` (‘-s 10 --maxseqs 1000’); this exceeds the sensitivity of the standard ‘sensitive’ MMseqs2 variant (`-s` 7.5), and is presented in order to demonstrate that the changes in sensitivity are due to `nail`’s pHMM alignment, not the choice of sensitivity flag. See Figure 4 for additional `nail` and MMseqs2 sensitivity flags. PSI-BLAST was run with default settings (provided a profile, with no iterative search). The resulting MMseqs2 and PSI-BLAST curves show a large sensitivity loss relative to pHMM annotation with HMMER, while `nail` recovers most of the difference.

As implemented, `nail` essentially re-scores candidate matches produced by MMseqs2, meaning that sequences filtered out by the MMseqs2 seed-finding stage can never be matched by the sparse FB algorithm of `nail`. To establish an upper bound on the recall gains possible with sparse FB, `nail` includes an option to compute the *entire* matrix for all candidates reported by MMseqs2 stage (‘--full-dp’). The corresponding curve is not shown here because it is essentially identical to that of the sparse implementation in `nail`. This demonstrates that loss of recall in `nail` relative to HMMER is due to limitations in candidates passing the initial filter, not failure of the sparse method, and highlights the value of future developments in fast candidate identification.

Note: this analysis accentuates the difference in real world performance of the tools because the benchmark is constructed to consist exclusively of hard-to-find matches. Furthermore, the performance gap may also be overstated due to the fact that the benchmark is built from Pfam sequences, which themselves were partly gathered using HMMER. Even so, the analyses agree with other observations of superior pHMM sensitivity [37, 23], and demonstrate that `nail` achieves most of the ideal pHMM sensitivity.

### nail approaches HMMER sensitivity at improved speed

Assessment of sequence annotation methods must consider the tradeoff between speed and sensitivity. In doing so, it is helpful to summarize the full recall curves from Figure 3 with a simple statistic. For each tool and setting, we compute the recall at 0.01 FP per search. Figure 4 presents a plot of run time (x-axis) versus this recall measure (y-axis) for annotation of the Pfam-based benchmark described above – an ideal tool will produce a point that is high (sensitive) and to the left (fast). The top half of the plot presents time/recall values for the profile based methods. HMMER’s hmmsearch is presented with default settings and shows the highest recall. PSI-BLAST is both slower and less sensitive than hmmsearch. Both `nail` and MMseqs2 are presented with several points corresponding to alternative selections of the mmseqs ‘sensitivity’ flag; when `nail` uses mmseqs as a prefilter with sensitive settings (`-s` 10 or `-s` 12), `nail` results are for default sensitivity settings (`-s` 10) and recovers most of the sensitivity gap between hmmsearch and mmseqs with speed similar to MMseqs2. DIAMOND with the `--ultra-sensitive` setting fits naturally on the MMseqs2 curve.

For “family pairwise” search involving only single-sequence queries, all tools produce much lower recall than with the profiles. `nail` and mmseqs show recall values similar to blastp at a ~4x speedup, and are nearly 10x faster than phmmer with a modest loss in recall.

We view these results as a conservative estimate of the speed benefits of the sparse Forward/Backward approach for full protein sequences, because the Pfam-based domain sequences are often quite short; the relative speed/recall tradeoff is expected to be increasingly in favor of sparse Forward/Backward for search involving longer sequences (see Figure 2A). We also tested with a variant of `nail` that implements calculation of the full DP matrix (not shown); this approach was much slower, but achieved the same recall as the fast variant, demonstrating that the speed boost achieved with sparse DP alignment results in essentially no loss in recall. Nearly all of the sensitivity difference between `nail` and HMMER3, for both profile and single sequence search, is the result of aggressive candidate filtering by the prefilter stage in MMseqs2, suggesting that an alternate ultra-fast alignment seed detection method is warranted.

### Sensitivity as a function of percent identity

Detecting the relationship between protein sequences becomes harder as their similarity decreases. To characterize this effect, we measured the rate at which recall declines as a function of query-target identity, and compared performance across tools. We extended the Pfam-based benchmark as follows.

#### Additional definitions

`pid` (q, t): the percent identity of (q, t) rounded to the nearest percent, as aligned in 𝔸 (the sum of exact column matches over total columns; columns with gaps in both sequences are ignored)

t → ψ: denotes that t is mapped to percent identity bin ψ

Beginning with the Pfam-derived benchmark from above:

- For each seed MSA 𝔸 in Pfam:
  - for each t_i_ ∈ 𝔸_T_:
    - let q^′^ = arg max pid(q, t_i_)
    - map t_i_ → pid(q, t_i_)
  retain the 10 target sequences mapped to the lowest percent identity bins (note: this ensures (i) we use the “hardest” available examples, and (ii) each domain contributes exactly 10 target sequences)
- The remaining target sequences from all Pfam domains comprise the set of true positives (note: the mapping of each t_i_ → ψ is maintained and written to a file)
- Use the same set of shuffled decoy sequences that were used in the previous experiment

We consider each individual target in conjunction with the set of decoys to be single *example*. The difficulty of the example is determined by the target’s percent identity to the most similar sequence among all sequences in the corresponding query MSA, 𝔸_Q_. This framework provides a scaffolding that allows us to evaluate the difference in sensitivity between sequence and profile queries at a reasonable approximation of the same level of difficulty (since the profile is built from sequences of at most ψ% identity to the target sequence).

One way that a sequence might be annotated is as the result of search with a profile learned from 𝔸_Q_. This process is modeled by the top lines in Figure 5, which show the recall of tools at a fixed false positive rate of 0.01 per query; this figure presents the relative efficacy of profile methods in hmmsearch, `nail`, mmseqs2, and psiblast (when run for a single iteration, given a profile query). These results show that all tools decay in sensitivity when the target is *<*25% identical to the best-matching sequence in 𝔸_Q_, and that `nail` maintains most of the sensitivity of HMMER.

**Fig. 5:**
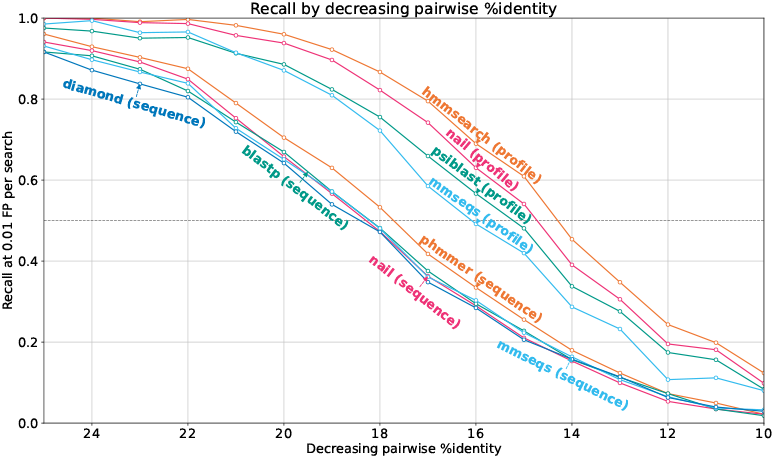
Recall vs. pairwise identity. The query/target pairs in the Pfam-based benchmark were binned by percent identity (rounded to the nearest percent) to demonstrate the loss of sensitivity as divergence between the query and target increases. `nail` variants were tested with `--mmseqs-s` `12.0 --mmseqs-max-seqs 2000`; MMseqs2 variants were tested with `-s 12.0 --max-seqs 2000 --seed-mode` full; DIAMOND was tested with `--ultra-sensitive`; HMMER (hmmsearch, phmmer) and BLAST (psiblast, blastp) were tested with default parameters.

An alternative way to annotate a target sequence is to search for it with each sequence from 𝔸_Q_ in turn, so called “family pairwise search”. Because we know in advance which sequence from 𝔸_Q_ has greatest identity to the target, we searched the target and all decoys with just that best matching query sequence; this is essentially an idealized family-pairwise search, which uses the likely best-aligning query sequence and minimizes false positive risk that would be introduced by using other query sequences. The lower lines of Figure 5 show that single sequence search is much less sensitive than profile-based search, and that all tools produce similar sensitivity (with phmmer yielding slightly greater sensitivity.

### Large-scale Pfam annotation of MGnify

To explore a real-world application that demands both sensitivity and speed, we evaluated the annotation of all ~2.4 billion proteins in the MGnify Proteins (2024 04 release [31]) with Pfam [26] pHMMs.

We restricted our analysis of MGnify annotation to `nail`, HMMER3, and MMseqs2. Since MGnify is not a curated database, we rely on coverage as a proxy for sensitivity. Because `nail` and HMMER share a scoring model, we could rely on the *trusted cutoff* scores found in the Pfam pHMMs when comparing the two in isolation, but MMseqs profiles do not follow the same scoring scheme. To bring analysis of MMseqs2 on equal footing with `nail` and HMMER, we devised a method to learn comparable per-family score cutoffs for all three tools.

For each domain 𝔸 in Pfam, we learn a score cutoff by establishing a large collection of decoy sequences. Rather than using shuffled sequences, we use reversed protein sequences. Reversed sequences almost certainly do not fold or behave the same as the original, yet they present traits that simple shuffled sequences do not and that are a common source of false matching between non-homologous sequence: short regions of low compositional complexity or repetitiveness that are often found in biological sequences. Importantly, sequences produce high scoring alignments to their reversals with surprising frequency, and can lead to overstatement of false match risk [17]; we take steps to overcome this.

#### Definitions

𝔸: a Pfam domain

μ: MGnify database (~2.4 billion sequences)

t_i_: the i^th^ sequence in MGnify

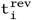: the reversal of sequence t_i_

`score` (𝔸, t): the score of aligning 𝔸 and t

E (ω): the E-value of score ω

𝔸 → ω: denotes 𝔸 is assigned cutof f score ω

To establish trusted cutoffs, we implement the following procedure:

- For each t_i_ ∈ μ
  - let 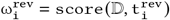
  - let ω_i_ = `score` (𝔸, t_i_) (note: most reversed sequences will not align to 𝔸; in these cases it is not necessary to compute ω_i_)
  - retain ω^rev^ if E (ω_i_) ≥ 1E-3
- all remaining ω^rev^ are considered scores of alignments to false positives
- assign 𝔸 → Ω_3_, where Ω_3_ is the third largest score among all ω^rev^; we refer to this assigned cutoff as *h*_*cut*_.

An analogous per-family cutoff *n*_*cut*_ is learned for `nail` by applying the same procedure to `nail` scores.

We randomly distributed the ~2.4 billion MGnify protein sequences into 1000 sets of ~2.4 million sequences each. This framework provides small, uniformly sized sets that can be easily batched and searched in parallel. We batched the data in this way to enable ideal search speed with HMMER3, which is known to scale poorly beyond ~4 threads on a single search [12, 5]; MMseqs2 and `nail` both scale well, and do not require small batching. Labeling all batches with Pfam required 7 hours with MMseqs2 (using sensitive settings; `-s` 7.5, `--max-seqs` 2000) and 73.9 hours using `nail` (with default `--mmseqs-s` 10.0 `--seed-mode` prog). With HMMER3 requiring 639.8 hours*, `nail` was 8.7x faster than HMMER and only 10.5x slower than MMseqs2. Table **??** shows times for other sensitivity settings of MMseqs2 and `nail`. While MMseqs2 remains faster than `nail` with default settings, its recall is lower; even extreme sensitivity settings cause it to be slower and still produce lower recall.

Many MGnify proteins are matched by more than one Pfam family above their respective thresholds, typically with one family dominating in score. To focus on hits that influence a target’s primary annotation, we apply a best-family filter that retains only the highest-scoring (family, target) pair per target. The filter operates separately on the hits for each tool.

Table 2 reports hit counts and recovery rates, grouped by HMMER’s reported domain structure. The first columns report hits for which HMMER recovers a single domain. The next columns report hits for which HMMER reports multiple domain hits, with at least one individual domain score exceeding the family-specific cutoff defined above. The final columns report multi-domain hits in which no single domain alignment clears the cutoff, meaning that HMMER’s call rests on summing several sub-threshold alignments.

**Table 2.** Hit counts (in thousands) for HMMER3, nail, and MMseqs2 on the MGnify database with Pfam profiles. Percentages are relative to the HMMER row. **Single-domain**: targets with exactly one positive-scoring domain alignment. **Multi**, ≥**1 dom** *> h*_*cut*_: targets with multiple HMMER domains, where at least one individual domain alignment exceeds the per-family score cutoff. **Multi, no dom** *> h*_*cut*_: targets with multiple HMMER domains, where no single domain alignment clears the threshold. Due to HMMER’s large runtime, its results were gathered on a cluster with little control over runtime; to report times, we performed search on 0.1% of MGnify, then extrapolated the full run time from this result.

| Tool | Single domain |  | Multiple domains |  |  |  | Run time |
| --- | --- | --- | --- | --- | --- | --- | --- |
| | Hits (K) | % | $\geq 1 \text{ domain} > h_{cut}$ | | no domain $> h_{cut}$ | | |
|  |  |  | Hits (K) | % | Hits (K) | % |  |
| HMMER | 1,939,265 | 100.0% | 178,356 | 100.0% | 148,839 | 100.0% | 639.8 hours* |
| nail ($s = 12$) | 1,862,426 | 96.0% | 171,806 | 96.3% | 25,161 | 16.9% | 133.1 hours |
| nail ($s = 10$, default) | 1,818,815 | 93.8% | 168,519 | 94.5% | 24,241 | 16.3% | 73.9 hours |
| nail ($s = 9$) | 1,741,385 | 89.8% | 160,837 | 90.2% | 22,497 | 15.1% | 51 hours |
| MMseqs2 ($s = 12$) | 1,451,342 | 74.8% | 131,805 | 73.9% | 23,601 | 15.9% | 86.7 hours |
| MMseqs2 ($s = 7.5$) | 1,351,872 | 69.7% | 115,475 | 64.7% | 18,228 | 12.2% | 7 hours |

**Table 3.**
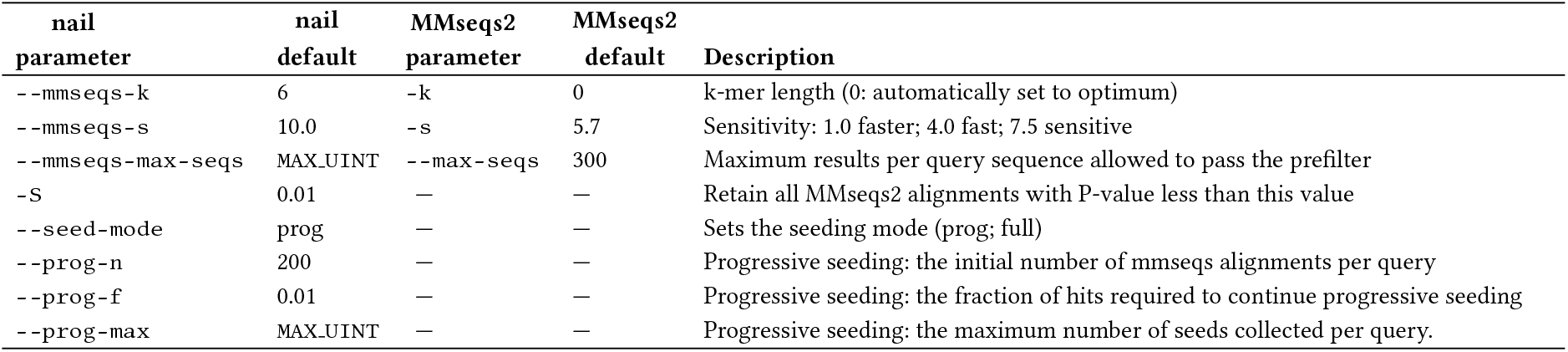
The nail command line parameters that impact seeding sensitivty, along with brief descriptions of their effects. For parameters that are directly passed through to MMseqs2, the MMseqs2 default is listed, and the description is copied from the `mmseqs search -h` command.

The first two groups are HMMER hits supported by at least one strong individual alignment. Across both, `nail` recovers 98% of HMMER’s hits and MMseqs2 recovers about 84%. Both `nail` and MMseqs2 recover a small fraction of the third group, when no single HMMER domain alignment alone clears threshold. The pattern reflects an architectural difference: HMMER’s full-sequence score is the sum of all per-domain contributions, including weak alignments, while `nail` and MMseqs2 each evaluate a single best-scoring alignment per (family, target) pair and do not aggregate sub-threshold evidence. The accumulated-score hits are examined in the following section.

### Many HMMER-only multi-domain hits accumulate score from weak, fragmentary matches

Within the best-family-filtered set, there are 4,601,210 (family, target) pairs for which HMMER’s full-sequence score exceeds *h*_*cut*_, HMMER reports more than one positive-scoring domain alignment, and neither `nail` nor MMseqs2 reports a hit. These are the pairs underlying most of the recovery gap in the rightmost columns of Table 2, driven by HMMER’s score-accumulation architecture (the J state, see [10]).

To understand what HMMER aggregates in these cases, we classified each pair by the spatial arrangement of its domain alignments on the HMM. Domain hit architectures are highly diverse and do not lend themselves to simple classification, but we have grouped matches into classes that provide a sense of the meaning and reliability of the matches (full algorithm in Supplementary Section 8). Categories are:

- *strong match*: at least one domain alignment alone exceeds *h*_*cut*_;
- *colinear match*: sub-threshold domains arranged co-linearly in both HMM and target space, such that the fragments can be chained together to approximately reach the family score cutoff (*h*_*cut*_);
- *shuffled match*: multiple sub-threshold domains are present that cannot be profitably arranged co-linearly in both HMM and target space, though their unordered sum is close to *h*_*cut*_ (these generally appear to be false positives);
- *pileup hits* in which overlapping copies pile up on the same pHMM region (no copy clears *h*_*cut*_ in any tier, so the target only shows weak similarity to the family in any single copy, but surpasses the cutoff because multiple copies are present; these are often false positives where score accumulates from repeated weak matches to a low-complexity or repetitive region of the target):
  - *pileup:full* (pileups span ≥70% of the pHMM): these may indicate that the target contains multiple weak copies of the family domain;
  - *pileup:fragment* (pileups span *<*50%): in our experience, these almost always appear to be false positives caused by accumulating score from many weak matches to a repetitive protein.
  - *pileup:mid* (pileups span 50% to 70%): these often appear to be false matches, but not always;
- *weak match*: matches that are not assigned to other classes above; in all cases, HMMER is only able to reach *h*_*cut*_ by accumulating scores from many low-scoring domains (*<*4 bits).

Figure 6 shows the category distribution by *n*_*pos*_, the number of positive-scoring domain alignments that HMMER reports for the pair, with a sample of 283,397 pairs (at most 50 per Pfam family). At *n*_*pos*_ = 2, *colinear match* dominates (38%): proteins with two partial copies of the domain in head-to-tail order, each below threshold but jointly above. At *n*_*pos*_ ≥ 3, *pileup:fragment* is the dominant class, with 57% at *n*_*pos*_ = 3, and rising to 73% at *n*_*pos*_ = 5; these consist of many overlapping copies of the same short region of the HMM, each contributing only a few bits to a total that crosses threshold. In 94% of *n*_*pos*_ ≥ 4 *pileup:fragment*, no individual domain alignment comes within 10 bits of *h*_*cut*_ (Supplementary Table S3).

**Fig. 6:**
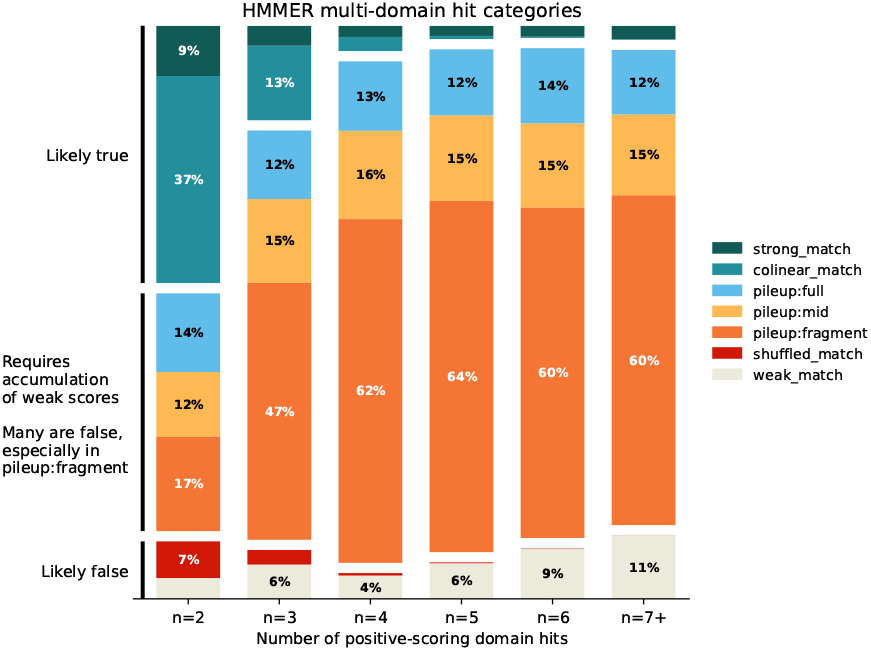
Domain-arrangement patterns in HMMER-only multi-domain hits. Each bar represents (family, target) pairs in the best-family-filtered set where HMMER’s full-sequence score exceeds *h*_*cut*_ but neither `nail` nor MMseqs2 reports a hit, grouped by *n*_*pos*_, the number of positive-scoring domain alignments HMMER reports for the pair. Category definitions are given in Supplementary Section 8. Each Pfam family contributes at most 50 (family, target) pairs (283,397 pairs total). At *n*_*pos*_ = 2, two co-linear sub-threshold copies (*colinear match*) is the dominant pattern; at *n*_*pos*_ ≥ 3, many overlapping copies of a fragment of the HMM (*pileup:fragment*) dominates.

The *pileup:fragment* pattern is consistent with repetitive or low-complexity regions of the target matching the same piece of the HMM many times. A genuine instance of the family domain would produce a single strong alignment to most of the HMM, not a pile of weak alignments to a fraction of it. Extrapolating sample fractions across the 4.6M HMMER-only multi-domain population, roughly 1.9M cases (~42%) fall in *pileup:fragment* or *weak match*; we would not confidently treat these as missed family instances. Another ~1.9M (~41%) fall in biologically credible classes (*strong match, colinear match, pileup:full*). Of these, the ~1.0M *colinear match* cases would be directly recovered by extending `nail` to report multiple alignments per (family, target) pair (see Discussion). Close to half of `nail`’s apparent multi-domain misses fall in *pileup:fragment* or *weak match*: score accumulated across many weak alignments to a fragment of the HMM, a pattern most readily explained by short low-complexity or repetitive regions of the target rather than by genuine domain homology. `nail`’s miss rate on credible HMMER multi-domain hits is much smaller than the gap in Table 2 suggests.

### Closed form for single-sequence query E-value parameter

`nail` inherits the significance estimation approach of HMMER3, which maps a Forward bit score *F* to an E-value *E* The E-value is the expected number of equal-or-better matches that would arise by chance in a database with no true homologs to the query (i.e. a measure of how surprising the match is). HMMER3 computes *P* (*F > t*) byt first applying an exponential tail to *F*:

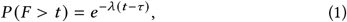

then adjusting for the size of the target database (*K* sequences) to produce *E* = *K* · *P* (*F > t*). As described in [10], the slope parameter *λ* for Eq. 1 can be computed with a closed form: for log-likelihood-ratio bit scores (log_2_), *λ* ≈ ln 2, independent of the model and the target (models with short length or low relative entropy require an edge effect adjustment, which HMMER3 implements as 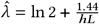, where where *L* is the model length and *h* is the average relative entropy per match state emission distribution). The location parameter *τ* has no analogous closed form, and is calibrated during model construction by aligning *n* = 200 i.i.d. random sequences of length *L* = 100 and fitting the resulting score distribution.

This simulation effort is generally a small part of the computation required to produce a pHMM from an MSA with `hmmbuild`. But when performing search with a single sequence query (as performed in `phmmer`, or optionally in `nail`), the time spent performing simulations to estimate *τ* can become a large fraction of the total run time. We observed that for single sequence queries, the *τ* learned through simulation is well-approximated by a simple closed form based on sequence length.

To establish the closed form estimate, we sampled sequences from the PANTHER 19.0 database, randomly selecting 1,500 families and concatenating their constituent sequences into a training set of 189,921 sequences. We also sampled a disjoint set of 1,500 PANTHER families to produce a held-out set of 193,872 sequences. For each sequence in both sets we ran `hmmbuild --singlemx` and recorded *τ* value (as the negated value from the pHMM `STATS LOCAL FORWARD` line) along with sequence length *L*. To quantify noise within the calibration itself, the training set was processed twice, once with the default random seed (`--seed 42`) and once with an alternate choice (`--seed 21`). To quantify noise between sequence sets, the held-out set was processed once with the default seed.

Ordinary least squares on *τ* values calibrated using `hmmbuild`’s default seed (42) on the the training set, yields the closed form estimate (restricted to two significant digits):

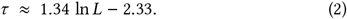

This estimate recovers training *τ* with Pearson *r* = 0.970 and mean |Δ*τ* | = 0.185 (Fig. 7a). The closed-form estimate is an equally good fit to the *τ* determined by `hmmbuild` on the training data with the alternate seed (seed = 21; Fig. 7b), and on the held-out set with default seed (Fig. 7c). A useful baseline is seed-to-seed noise: two `hmmbuild` calibrations of the training sequences with different random number seeds agree at *r* = 0.962, mean |Δ*τ* | = 0.210 (Fig. 7d). The closed form’s deviation from any single calibration run (Fig. 7a–c) is smaller than the deviation between two independent calibration runs.

**Fig. 7:**
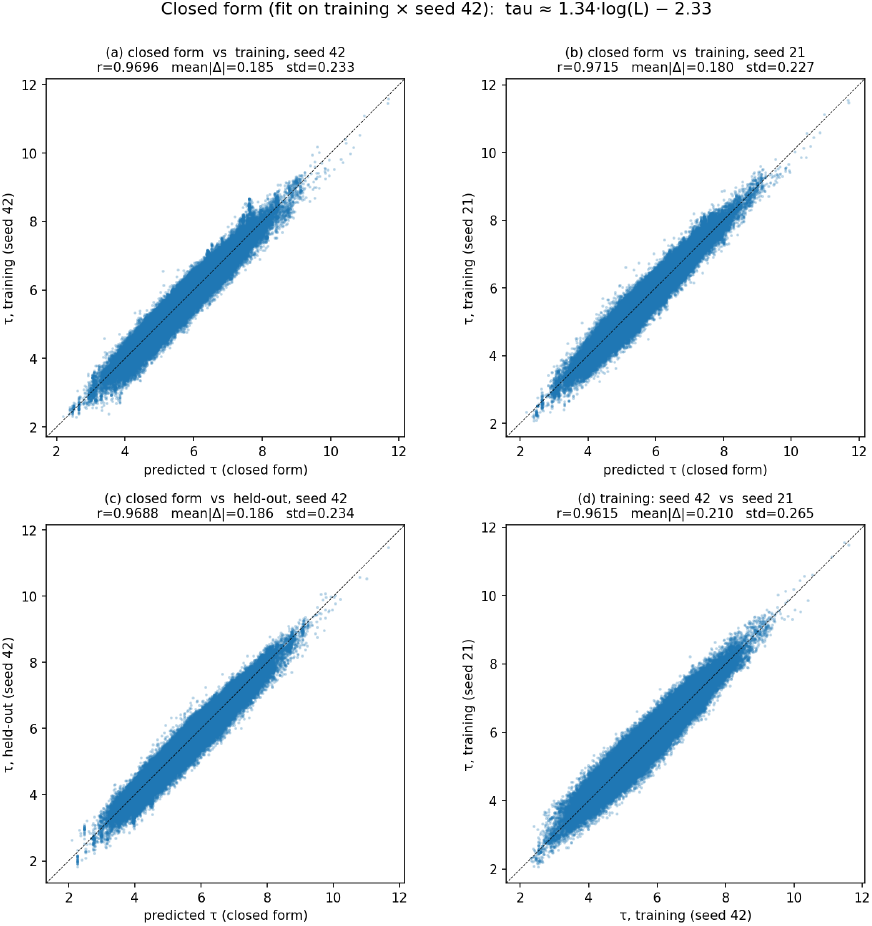
Closed-form *τ* prediction is statistically indistinguishable from re-running the calibration simulation. Each panel is a scatter of ~190,000 single-sequence HMMs built with `hmmbuild --singlemx`; axes are *τ* in bits with the unity diagonal drawn for reference. **(a)** Closed-form estimate (Eq. 2) vs. calibrated *τ* on the training set with default `--seed 42` (*r* = 0.970, mean |Δ*τ* | = 0.185). **(b)** Same estimate vs. the training set re-calibrated with `--seed 21` (*r* = 0.972, mean |Δ*τ* | = 0.180), showing that the fit (trained on seed 42) generalizes to a differently seeded calibration of the same sequences. **(c)** Closed-form estimate vs. the disjoint held-out set with default seed (*r* = 0.969, mean |Δ*τ* | = 0.186), showing generalization to unseen sequences. **(d)** Training *τ* at `--seed 42` vs. `--seed 21` (*r* = 0.962, mean |Δ*τ* | = 0.213), establishing the seed-to-seed noise floor of the calibration itself. The point-cloud width and residual statistics in (b) and (c) match those in (d): the closed form introduces no more error than a different random number generator seed already would.

Note that |Δ*τ* | = 0.210 translates to a negligible 1.16-fold change in E-value, and the largest observed prediction error over all comparisons in Figure 7 (1.507) translates to only a 2.83-fold change in E-value.

Using this closed form computation removes the need to run the calibration simulation, improving `nail` speed for search with single sequence queries. For pHMMs built from multiple sequence alignments, `nail` continues to rely on `hmmbuild` for both model construction and *τ* calibration. We explored an analogous closed-form fit in the MSA regime, but found the correlation to be slightly weaker in general, and more chaotic for short pHMMs; we expect that a closed form may be possible, but leave its derivation for future work.

## Methods

The `nail` pipeline consists of the following steps:

- For each query profile (or sequence), rapidly prefilter target sequences to identify good candidates full sequence alignment;
- For each surviving query-target pair:
  - identify alignment matrix positions to serve as seeds for sparse matrix computation;
  - perform profile-to-sequence alignment using a new sparse implementation of the Forward/Backward algorithm;
  - perform supplementary analyses such as composition bias correction;
- Report pairs with significant Forward/Backward scores, producing alignments and statistics as requested

We begin the description of the `nail` implementation by focusing on the sparse Forward/Backward implementation, as this is the primary novelty of `nail`. This is followed by details of the prefiltering process, how this connects with sparse Forward/Backward, and supplementary aspects of the `nail` pipeline.

### Profile HMMs: preliminaries

Profile HMMs (pHMMs) resemble position-specific scoring matrices (PSSMs) in capturing per-position properties of a sequence family, but specify transition and emission probabilities rather than log-probability-ratio scores, and are aligned via probabilistic inference algorithms rather than classical Smith-Waterman-style algorithms.

#### Definitions

- *A*: an alphabet
- *B*: the background frequencies of amino acids
- *Q*: a length-*m* query pHMM {*q*_1_, *q*_2_, … *q*_m_}, where each position *q*_j_ is represented by a tuple of states (*M*_j_, *I*_j_, *D*_j_)
- *Q*_j_: the prefix of *Q* up to position *j*; {*q*_1_, *q*_2_, …, *q*_j_}
- *T*: a length-*n* target sequence {*t*_1_, *t*_2_, …, *t*_n_}, where each character *t*_i_ ∈ *A*
- *T*_i_: the prefix of *T* up to position *i*; {*t*_1_, *t*_2_, …, *t*_i_}.
- *S* → *S*^′^: denotes a transition from state *S* to state *S*^′^ where *S, S*^′^ ∈ {*M, I, D, N*, *C*}
- *π*: a path through the states of *Q*
- Π_T|Q_: the set of paths *π* through *Q* capable of emitting *T*
- *P* (*T* |*π*): the probability of the path *π* emitting the target sequence *T*

### States & emissions

A profile HMM *Q* (see Figure 8) is made up of a collection of states that emit characters from an alphabet *A* (here, the amino acid alphabet), built from a multiple sequence alignment (MSA). Each mostly-occupied column *j* in the MSA produces a tuple (*M*_j_, *I*_j_, *D*_j_), representing position *q*_j_ in *Q*, containing:

**Fig. 8:**
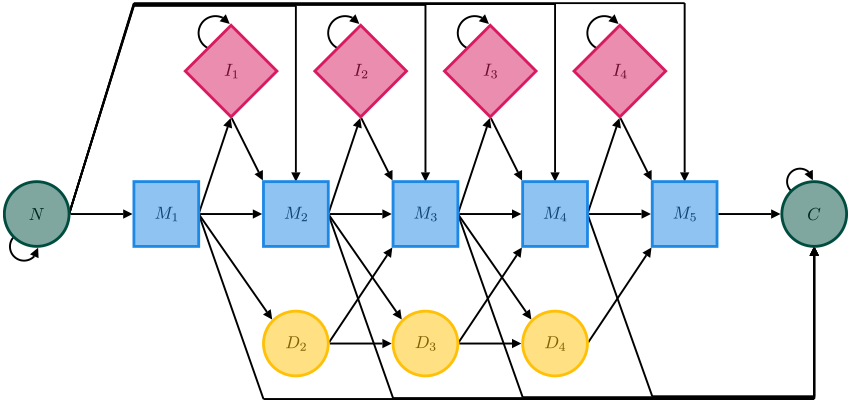
The profile HMM architecture, shown here with a 5-position core. Match states *M*_*j*_ (blue squares) emit residues with position-specific distributions learned from a family multiple sequence alignment; insert states *I* _*j*_ (pink diamonds) and the flanking background states *N*, *C* (green circles) emit from the background distribution *B*; delete states *D* _*j*_ (yellow circles) are silent. Position-specific transitions govern the choice of match, insertion, or deletion at each core position. Transitions *N* → *M*_*j*_ and *M*_*j*_ → *C* allow alignment to begin and end at any core position, with unrelated flanking residues handled by the background states. The architecture is generative: it produces sequences consisting of a family homolog optionally flanked by unrelated residues. Assuming an observed target *T* was generated this way supports inference of *T*’s family membership.

- A match state (*M*_j_), which emits from a distribution based on the contents of j^th^ MSA column [35]; these states model alignment of a character *t*_i_ from *T* to a position *q*_j_ in *Q* (note: the match emission distributions are analogous to the scores in a PSSM)
- An insert state (*I*_j_), which emits from a distribution that in HMMER is identical to the background prior *B*; these states are used to model insertions relative to *Q* (note: in principle, position specific insert emissions may be used)
- A delete state (*D*_j_), which is silent (no emission); these states are used to model deletions relative to *Q*

We refer to the set of all *m* tuples as the *core model*. There are two structural exceptions in the core model: (i) the first position *q*_1_ is represented as (*M*_1_, *I*_1_, –), and (ii) the last position *q*_m_ is represented as (*M*_m_, –, –). The core model is supplemented by two flanking *background* states (*N*, *C*), which both emit from *B*, and are designed to account for all target residues not aligned to positions in the core model.

Note: *Q* may also be a single sequence; in this case, a pHMM is built with (i) uniform insertion/deletion transition probabilities and (ii) emission distributions derived from the probabilities used to produce a classical SW-style substitution matrix (e.g. BLOSUM62).

### Transitions & paths

In the following sections, we will refer to a path *π* through the states of *Q*, which has the following properties:

- Begin in an implicit *source* state, then either transition to:
  i. the core model in some *M*_j_, or
  ii. the background *N* state
- Upon entering the *N* state, either transition to:
  i. the core model at some match state *M*_j_, or
  ii. the *N* state (self-loop)
- Upon entering the *M*_j_, either:
  i. remain in the core model via the transition scheme:
    - *M*_j_ → *M*_j+1_
    - *M*_j_ → *I*_j_
    - *M*_j_ → *D*_j+1_
    - *I*_j_ → *M*_j+1_
    - *I*_j_ → *I*_j_
    - *D*_j_ → *M*_j+1_
    - *D*_j_ → *D*_j+1_
  ii. or, escape by transitioning to either:
    - the background *C* state, or
    - the implicit *terminal* state
- Upon entering the *C* state, either transition to:
  i. the *C* state (self-loop), or
  ii. the implicit *terminal* state

In this way, a path *π* represents sequences that include a variable length homolog of *Q*, optionally flanked on either side by unrelated background sequence; the set of characters emitted by the states in *π* produce a sequence of this form. Notably, the probabilities of transitioning to insert and delete states are informed by the source MSA, and they directly correspond to position-specific tolerance of insertions and deletions during alignment.

### Hidden states & inference

Given*T* and a path *π*, *P* (*T* |*π*) is the product of the corresponding transition and emission probabilities. In inference, we observe target sequence *T* and assume that *Q* emitted it (i.e. that *Q* and*T* are homologous), but *π* is *hidden*. To make a statement about whether *Q* and*T* are related, we select some *π* a posteriori. Among the many valid paths in Π_T|Q_, a natural choice is to select the most probable path:

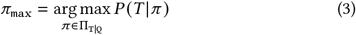

We can compute *P* (*T* |*π*_max_) with dynamic programming via the Viterbi algorithm [39]. We omit the recurrence here, as Viterbi is not computed by `nail` and differs from the Forward recurrence (described in the following section) only by replacing the *sum* function with a *max* function. The implied *π*_max_ is recoverable via traceback through the dynamic programming matrices; this is essentially equivalent to the scheme used in Smith-Waterman, BLAST, MMseqs2, DIAMOND, and others [8, 15]. The states along *π*_max_ represent the most probable alignment between *Q* and *T*:

- any *t* ∈ *T* emitted by a background state *N* or *C* is excluded from the alignment;
- *t*_i_ is aligned to *q*_j_ when *M*_j_ emits *t*_i_;
- *t*_i_ is aligned to a gap between *q*_j_ and *q*_j+1_ when *I*_j_ emits *t*_i_ (an insertion relative to *Q*);
- *q*_j_ is aligned to a gap between *t*_i_ and *t*_i+1_ when *π*_max_ visits *D*_j_ (a silent deletion relative to *Q*)

We score the Viterbi alignment as a log likelihood ratio:

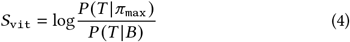

where *P* (*T* |*π*_max_) is the product of the transition and emission probabilities along *π*_max_, and 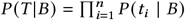 is the probability of *T* under the background model (each residue independently drawn from *B*). The *D* state contributes no emission, and because the *I, N*, *C* states emit from *B*, their emission factors cancel in the ratio; indels and flanking sequence are thus ‘penalized’ only by transition costs.

### The Forward algorithm

As divergence between sequences increases, their relationship becomes more difficult to identify. In these cases, we can instead consider each *π* ∈ Π_T|Q_ as evidence in support of the relationship between *Q* and *T*. In other words, we take the sum of the probabilities of all paths:

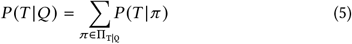

The probability sum *P* (*T* |*Q*) is computed by the Forward algorithm, which is a small modification to the Viterbi algorithm. First, consider the paths 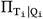 through *Q*_j_ = {*q*_1_, *q*_2_, …, *q*_j_} that are capable of emitting the subsequence *T*_i_ = {*t*_1_, *t*_2_, …, *t*_i_}. Note that, each 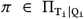 is a path *π*_S_ terminating in state *S* ∈ {*M*_j_, *I*_j_, *D*_j_, *N*, *C*}. Given a length *m* query pHMM *Q* and a length *n* target sequence *T*, Forward fills in:

i. three (*n* + 1) by (*m* + 1) matrices, *F* ^M^, *F* ^I^, and *F* ^D^, where:
  - 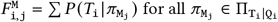
  - 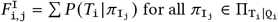
  - 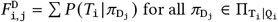
ii. two length-(*n* + 1) vectors, *F* ^N^ and *F* ^C^, where:
  - 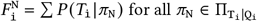
  - 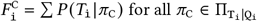

After initializing 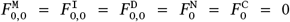, the remaining matrix cells are computed via the recurrence equations in Figure 9A. Once complete, we have:

**Fig. 9:**
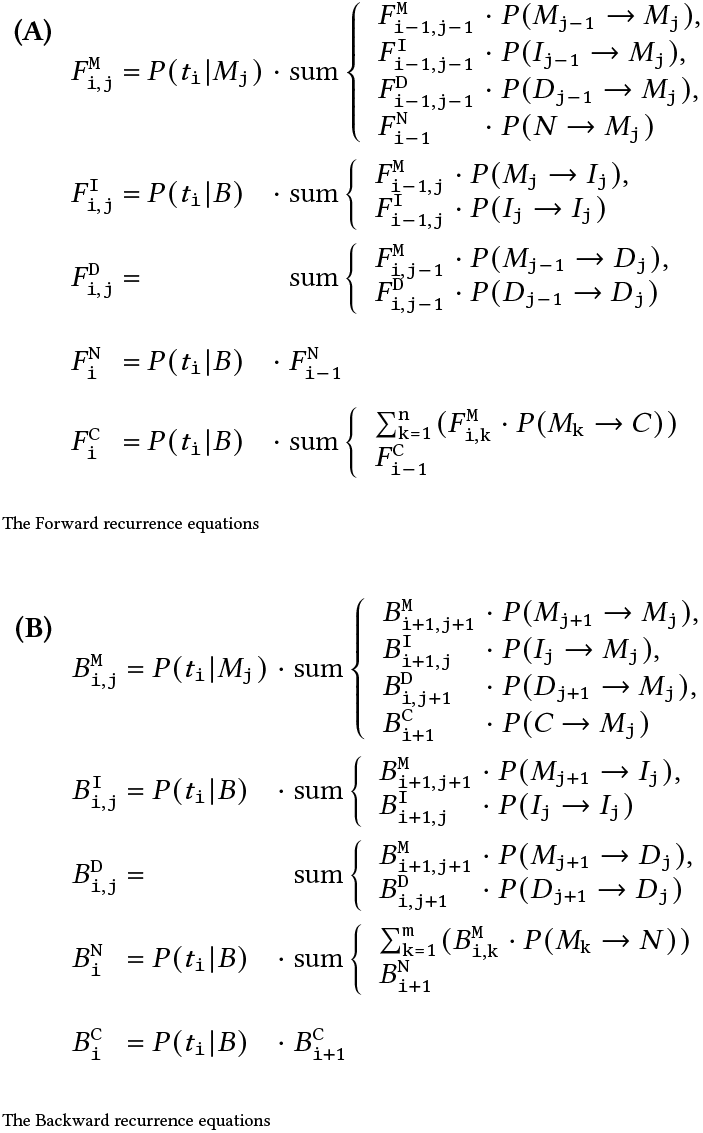
Forward (A) and Backward (B) recurrence equations for profile HMMs. Despite their apparently recursive form, the matrices can be computed in quadratic time by *dynamic programming*, filling a table in an order that ensures that any value is computed only after its dependencies have been resolved. Row-major order (upper-left to lower-right) is the most common fill pattern, but the dependencies also permit other orders such as sequential anti-diagonals [32], as is done in nail. Both recurrences chain products of probabilities, and can suffer from numerical underflow if computed directly. The Viterbi (max) recurrence sidesteps this by working entirely in log space, but the sums in Forward and Backward preclude that approach. To avoid moving values in and out of log space, nail instead uses a fast approximation of log(*P*_1_ + *P*_2_). Further acceleration is possible by scaling values directly to avoid log conversion altogether [11].

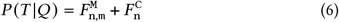

Then, we can assign a score as:

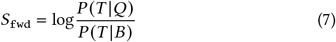

### Backward

The Forward algorithm produces a homology score *S*_fwd_, but does not directly produce an alignment between *T* and *Q*. For that, we also need the analogous Backward algorithm, which mirrors Forward by traversing *T* in reverse and inverting the pHMM transitions. The corresponding recurrence is shown in Figure 9B.

### Posterior probabilities, alignment boundaries, alignment

The Forward and Backward matrices can be combined to produce posterior probabilities for each cell in the alignment matrix [8], which in turn serve as the basis for maximum expected accuracy alignment [20].

After application of the recurrence in Figure 9, the matrices *F* and *B* store Forward and Backward probability sums associated with the states *M, I, D, N*, *C* for each target residue *t*_i_. The product of corresponding

Forward and Backward values (equations 8, 9) produces a matrix *Z* of joint probability sums. Each entry 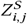 is the sum, over all alignments between *Q* and *T* that have state *S* _*j*_ emit *t*_*i*_, of the probability of those alignments:

- 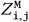 is the joint probability sum over all alignments that align the residue *t*_i_ to the profile position *q*_j_
- 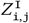 is the joint probability sum over all alignments that align the residue *t*_i_ to a gap following position *j* in *Q*
- 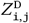 is the joint probability sum over all alignments that align the model position *q*_j_ to a gap following position *i* in *T*.

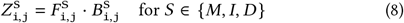

For the model’s background states (*N*, *C*), each value 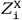 is the analogous joint probability sum over all alignments that emit *t*_i_ from background state *X* (i.e., alignments that exclude *t*_i_ from the homologous region).

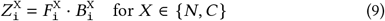

These joint probability sums can be normalized to obtain posterior probabilities 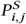 that target residue *i* is associated with query model position *j* via state *S* in the correct alignment [8].

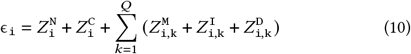

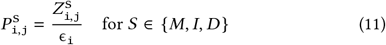

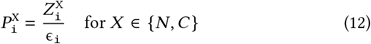

These posterior probabilities can then be used to identify a maximum expected accuracy (MEA) alignment [20], computed by a simple dynamic programming algorithm that maximizes the sum of posterior probabilities across aligned positions. The MEA alignment provides both the specific boundaries of the alignment and the alignment itself.

### Efficient search for high-probability cloud in Forward/Backward matrices

The Forward/Backward computation captures the total probability of all possible alignments, but does so by filling matrices of quadratic size (the product of the lengths of *T* and *Q*). Most paths through these matrices carry so little probability that excluding them has negligible effect on the result (Figure 1); `nail` exploits this by identifying a sparse cloud of high-probability cells and limiting subsequent computation to that cloud.

The cloud-identification method, which we call *Cloud Search*, resembles the X-drop algorithm used in maximum-score alignment tools such as BLAST [1]. Starting from a seed that contains likely high-probability cells, `nail` expands forward and backward across the matrices to grow a cloud that captures essentially all relevant probability mass. This cloud is the basis for all downstream analysis.

### Cloud Search by pruned anti-diagonal completion

The method proceeds as follows

- Cloud Search is initiated with a pair of alignment matrix cells, *begin* and *end*. As currently implemented, this pair is taken from an MMseqs2 alignment between *Q* and *T* (Figure 10A): the first and last positions of the alignment specify the *begin* cell (*i*_b_, *j*_b_) and *end* cell (*i*_e_, *j*_e_). (note: in principle, a cell pair could be produced by some other seed-finding approach)
- Cloud Search flood-fills the matrices forward (down and right) from the *begin* cell, extending until pruning conditions are met – Figure 10B.

**Fig. 10:**
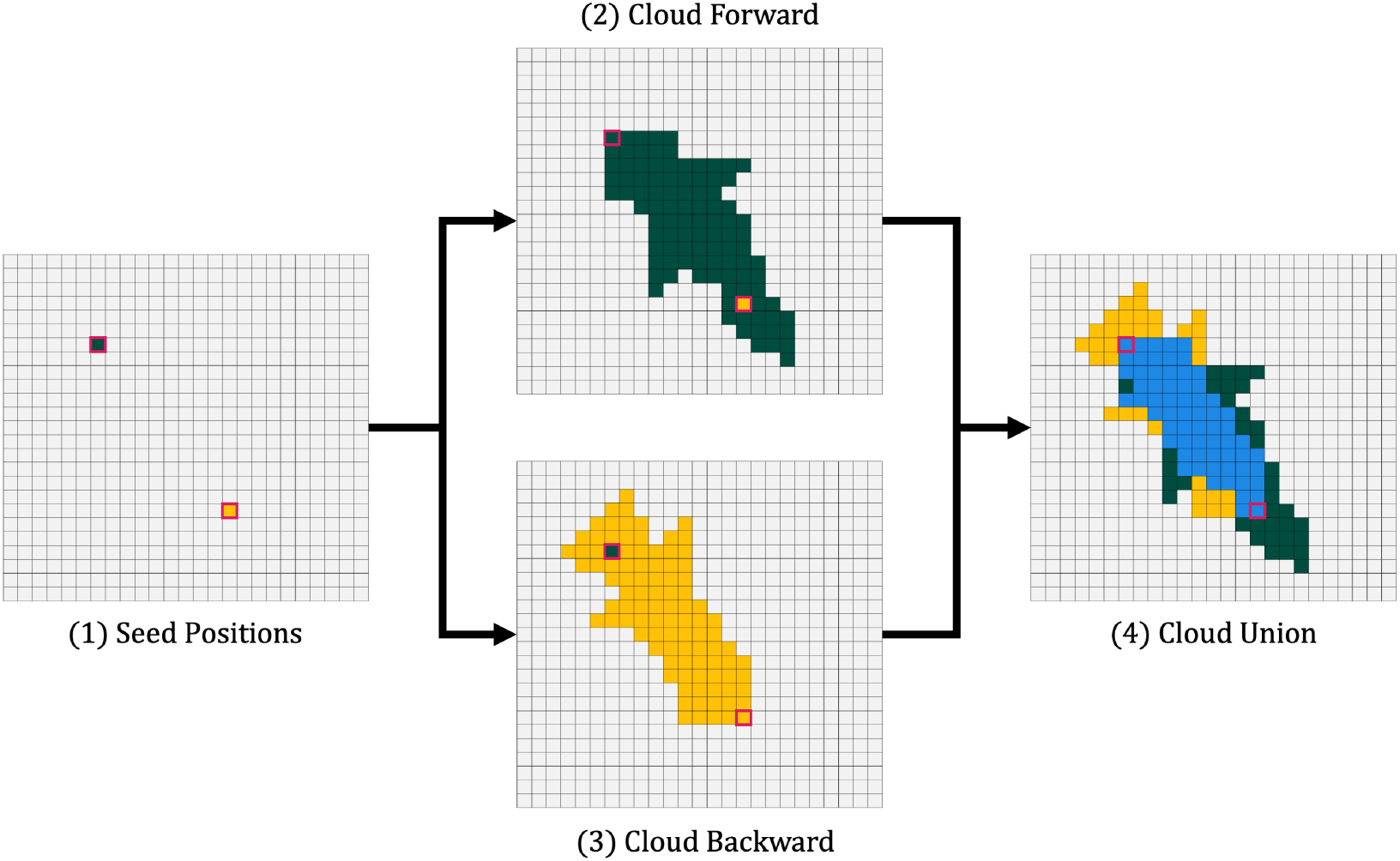
Cloud Search. In this schematic representation of Cloud Search: (A) An alignment from MMseqs2 is used as the source of begin- and end-points (green and yellow; these could come from any source). (B) Calculation is performed in the forward direction (moving down and to the right) from the begin point by filling in one anti-diagonal at a time, pruning each diagonal in from the ends based on score-dropoff conditions; this typically extends beyond the provided end point. (C) A similar flood fill pass is performed in the reverse direction starting from the provided end point, moving up and to the left. (D) The union of the two resulting spaces is identified as the sparse cloud.

After initializing 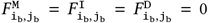 (green cell in the upper left), neighboring cells down and right of (*i*_b_, *j*_b_) are computed in anti-diagonal fashion, first filling the two cells (*i*_b+1_, *j*_b_) and (*i*_b_, *j*_b+1_), then the three cells below those, and so on. Based on the recurrence, each cell on one anti-diagonal pushes values to recipient cells in two subsequent anti-diagonals; because of this push-based transfer of information, the only cells touched on one anti-diagonal are those that are reachable from some active cell on the previous two anti-diagonals. Beginning from (*i*_b_ *j*_b_), all reachable anti-diagonal cells are computed and retained, until the search has advanced *γ* anti-diagonals from the start (default: 5). After this, when an anti-diagonal has been computed, two pruning conditions are applied to constrain expansion of the search space.

- Once all possible cells in an anti-diagonal *d* have been computed, the maximum value for that anti-diagonal is captured as max_*d*_. Cells are retained only if their score is within *α* of max_d_; others are pruned. Scores at this point are captured in *nats* (natural logarithms), with default *α* = 12, so that this effectively prunes cells on an anti-diagonal that have probability that is ~160,000-fold lower than the most-probable cell on that anti-diagonal.
- As flood fill continues, the overall best-seen score across all computed anti-diagonals is captured as max_*o*_. After an anti-diagonal has been computed, cells at its ends are pruned while their scores remain below the combined threshold max(max_*d*_ − *α*, max_*o*_ − *β*). With a default *β* = 20, the second term of this combined threshold acts like an X-drop [41] criterion, pruning cells with ~500 million-fold reduction from the best seen overall value. When all cells in an anti-diagonal are pruned, the flood fill stops.

Pruning is performed using the best score among the Match, Insert, and Delete states for each cell, and all scores are maintained in log space. The result of this phase is a set of cells expanding down and right from (*i*_b_, *j*_b_), schematically represented as dark green cells in Figure 10B. This cloud of cells typically remains in a tight band around the maximum-probability (Viterbi) path, though it can expand in cases where a pHMM has a very high insertion probability. Importantly, this cloud search phase typically extends well beyond the initial *end* cell (*i*_e_, *j*_e_), meaning that a conservative choice of initial points does not constrain the Forward cloud search.

- After the Forward Cloud Search phase, a similar Backward pass is performed, beginning at (*i*_e_, *j*_e_) and flood-filling as in the previous stage, but up and to the left (Figure 10C; yellow cells).
- Cloud Search concludes by taking the union of the Forward and Backward clouds (Figure 10D). This establishes a set of cells representing a non-negligible expansion around the range bounded by the initiating cells (*i*_b_, *j*_b_) and (*i*_e_, *j*_e_)

### Linear space requirement for computing Cloud Search

The forward and reverse passes of Cloud Search can be computed in linear space, using a (3 · *m*) matrix, in which each row holds the dynamic programming values computed along one anti-diagonal. In general, the *n*^*th*^ anti-diagonal, *d*_*n*_, is assigned to row (*n* mod 3). For a given matrix cell 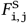, its anti-diagonal is given by *d*_*n*_ = *i* + *j*. Then, the value is stored in the cloud matrix at row ((*i* + *j*) mod 3), column *j*. Modifications to the recurrence equations follow naturally.

During the forward pass of Cloud Search, the values computed along anti-diagonal *d*_n_ depend on the values computed along the previous anti-diagonals *d*_n−1_ and *d*_n−2_. The cloud matrix access pattern satisfies those dependencies: the values along *d*_n_ are stored in the row that previously contained the (now retired) values of *d*_n−3_, while the previously computed values of *d*_n−1_ and *d*_n−2_ remain available. Similarly, during the reverse pass, the values along *d*_n_ are stored in the row previously containing the values of *d*_n+3_ with the values on *d*_n+1_ and *d*_n+2_ retained. Figure 11 gives an example of the cloud matrix access pattern during a forward pass of Cloud Search.

**Fig. 11:**
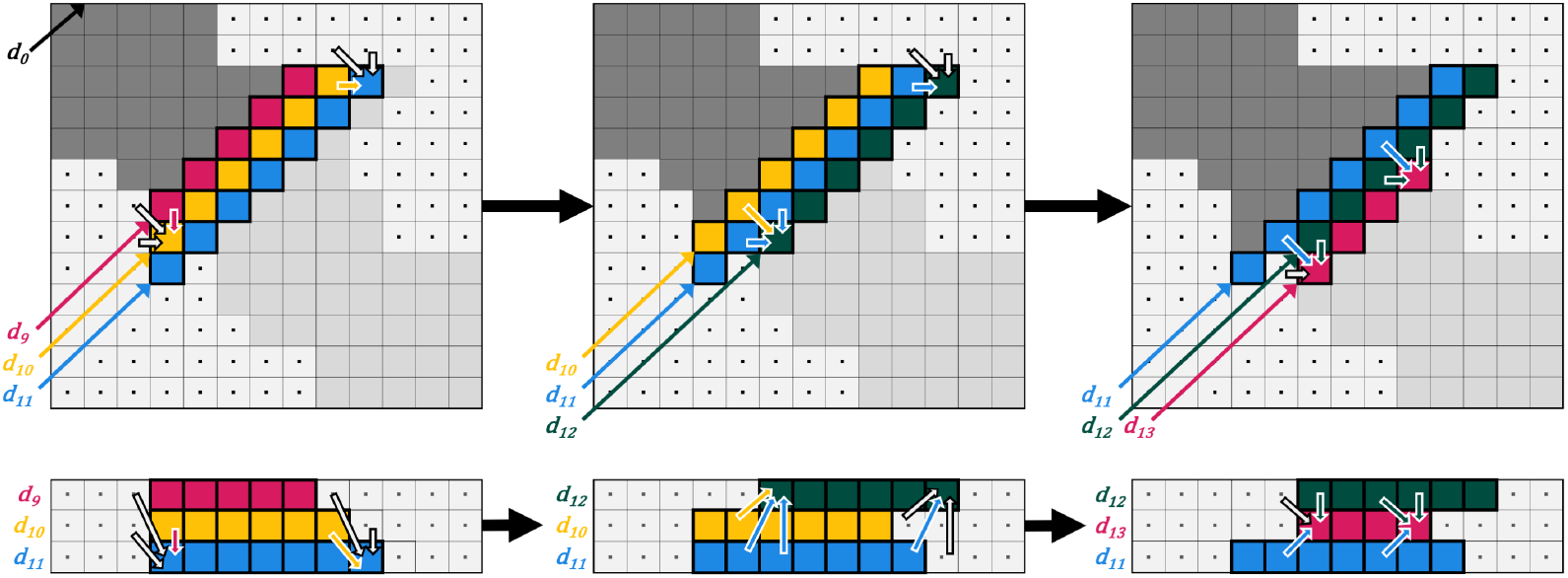
Example anti-diagonal access pattern. This example shows the implicit DP matrix (top) and the state of the cloud matrix (bottom) when anti-diagonals *d*_11_ (left example), *d*_12_ (middle example), and *d*_13_ (right example) are being filled during the forward pass of Cloud Search. In both representations, the data dependency patterns are shown with arrows. Note that in the implicit DP matrix, the dependencies follow the same patterns at each step, but, in the cloud matrix, the relative positioning (in memory) of the dependencies changes.

Once an anti-diagonal has been computed and pruned, the positions (in the implicit complete matrix) of its lower left and upper right cells are stored; these two cells describe an anti-diagonal cloud bound: each cell along the anti-diagonal between the two bounding cells is included in the sparse cloud. In this fashion, the cloud bounds are stored in linear space, with at most 4 · (*m* + *n* − 1) values (two (*i, j*) pairs per anti-diagonal) describing a cloud that spans the full width and height of the complete

matrix.

### Cloud union

After completing the forward and reverse passes of Cloud Search, the union of the two clouds is taken, as shown in Figure 10D.

It is possible for the clouds identified by the Forward and Backward passes to not intersect, typically due to a long gap spanning a region of very low alignment quality. In such cases, `nail` attempts to recover by iteratively re-running cloud search until the clouds intersect; at each iteration the pruning parameters (*α, β, γ*) are scaled to be more permissive. The maximum number of attempts and scaling factor configurable via the command line (`-a`, default 5; `-f`, default: 0.5). When the iteration threshold is reached, `nail` discards the search clouds and defaults to filling in a rectangular DP matrix bounded by the start and end seed positions.

### Sparse Forward/Backward to recover score and alignment

With a cloud of non-negligible alignment matrix cells in hand, `nail` computes an approximation of the full Forward/Backward algorithm by filling in only cells within the cloud, implicitly treating all other cells as zero. The implementation achieves this using compressed arrays that correspond to the shape of the sparse cloud and by computing two length-*n* vectors of probabilities associated with the background states (N, C). The compressed arrays are organized as sets of values that correspond to each cloud-bounded anti-diagonal, with padding cells in between each set (e.g. [*d*_4_, −, *d*_5_, −, *d*_6_, −, *d*_7_]). When filling in the sparse matrices, pad cell values are set to zero, and other cells are computed based on the standard recurrence equations, with retrieval of data via logical row and column indices.

### Bias correction

Alignment scores are computed as a ratio between the probability of observing a sequence under a model of homology (i.e. assuming the target sequence is related to the query) and a background model (i.e. the probability of observing the target sequence by chance). When both model and sequence show substantial and similar composition bias, large scores can result entirely due to chance alignment of nonhomologous residues. This results in inappropriate estimates of the significance of relationship (E-value) between the two sequences.

Most modern tools correct for this by adjusting scores in cases of biased composition, approximating what the score would have been under a more appropriate background. For protein search, HMMER3 does this by computing an expected score for each residue *c* in the alphabet *A* under the conditions of the alignment of *Q* to *T*. We adapted HMMER3’s method to accommodate `nail`’s sparse F/B alignment matrices, as described below; all sums in the procedure run only over cells inside the sparse cloud, with positions outside contributing zero:

- Compute vectors σ and ζ, where σ_j_ is the probability that the match state *M*_j_ emits a residue, and ζ_j_ is the analogous quantity for the insert state *I*_j_ These values are determined from the column-sums of the posterior probability matrices *P* (see equation 11). Positions outside the sparse cloud contribute no value to the sum; within the cloud, most σ_j_ values are ≈ 1, but a few positions at the edges and around gaps will be *<* 1:

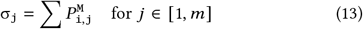

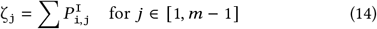
- Compute a vector ϕ where each element ϕ_i_ is the posterior probability that the corresponding target residue *t*_i_ was emitted by one of *Q*’s core states (the probability that *t*_i_ is included in the Maximum Expected Accuracy alignment, whose span in *T* we denote [*a, b*]). Target positions outside the cloud are implicitly assigned a value ϕ_i_ = 0:

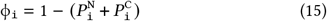
- Compute a vector χ where each element χ_c_ is the expected likelihood ratio of model-emitted to background-emitted residue *c* normalized by the expected alignment length:

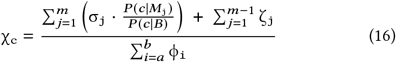

The numerator has two parts: the first sum captures the expected likelihood-ratio contribution from match emissions for residue *c*; the second sum captures the expected total insert-state usage, which is independent of *c* because `nail` follows HMMER3’s convention of setting insert emissions equal to the background. The denominator is the expected length of the alignment. When χ_c_ *>* 1, the aligned region of the model tends to emit *c* at a rate above background (i.e. the aligned model and target are similarly biased, and the score should be corrected).
- For each target residue *t*_i_ within the cloud, the expected score for that residue is computed by taking the log of 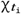 weighted by the core state posterior probability ϕ_i_. Effectively, this sets the influence that each residue has on the bias score in proportion to the confidence that it is included in the alignment. The sum of these weighted log ratios is then mixed with a strong prior against composition bias (weight ω = 1/256, following HMMER3) to produce the composition-bias score adjustment θ (equation 17).

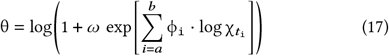

The data-driven sum 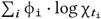 only dominates when the evidence for bias is large. As a consequence, θ ≥ 0 always, so the bias correction can only reduce a raw alignment score, never inflate it.

### Forward filter

Though sparse Forward/Backward is fast, many input alignment candidates will produce such a low-quality alignment that they will not end up being reported. To avoid time spent analyzing such candidates, `nail` computes a P-value from the sparse Forward score and discards candidates with P-value *>*1e-4, so only 0.01% of unrelated sequences are expected to pass.

### Identifying candidates for Forward alignment

As described earlier, an initial seeding step is required in order to identify a collection of query-target pairs as candidates for cloud search, along with anchors for the Cloud Search process. The `nail` pipeline currently identifies candidates and corresponding anchors with the aid of two MMseqs2 modules [37].

First, `nail` runs the MMseqs2 **prefilter** module with permissive sensitivity settings (nail default: 10.0; MMseqs2 default: 5.7; MMseqs2 suggested for sensitive search: 7.5). Internally, this identifies seeds by finding co-diagonal pairs of good-scoring k-mer matches, then computes a vectorized ungapped alignment along each seeded diagonal and returns the maximal score for the query/target pair.

Next, `nail` runs the MMseqs2 **align** module, which computes an optimal gapped Smith-Waterman alignment for each pair surviving the prefilter, accelerated by Farrar’s striped SIMD scheme [13]. By convention, **align** filters results using E-values (mmseqs; **-e** parameter; MMseqs2 default: 1`E-3`), the expected number of equal-or-better matches that would arise by chance in a database with no true homologs. So that the filter behaves consistently regardless of search space size, `nail` converts MMseqs2 E-values to (approximate) P-values, which measure alignment significance independent of search space conditions. To this end, a chosen P-value, *P*_seed_ (adjustable via `nail` flag **-S**), is converted to a comparable E-value target, *E*_align_ for MMseqs2, with

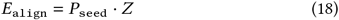

where *Z* is the target database sequence count (**note:** this is an approximate conversion due to differences between the Karlin-Altshul statistics used in MMseqs2 and the HMMER3 model). Then, *E*_align_ is used as the E-value cutoff parameter in the call to **align**. By default, `nail` filters out alignment candidates with P-value *>* 1`E-3`, meaning that 0.1% of non-homologous query-target pairs are expected to pass the filter. This filtering strategy this is similar to the Viterbi stage filter in HMMER3 and permits `nail` to search for distant matches that MMseqs2 **align** fails to report as significant.

### MMseqs2 --max-seqs and progressive seeding

The MMseqs2 **prefilter**’s sensitivity parameter (**-s**) has the most straightforward impact on the tool’s runtime and sensitivity characteristics. However, we have found that tuning the MMseqs2 `--max-seqs` parameter is challenging and vital, particularly as the size of search increases. Following the initial paired k-mer and ungapped alignment prefilter steps, MMseqs2 produces a list of candidates, sorted by their ungapped alignment (prefilter) score. The `--max-seqs` setting determines the number of prefiltered results that are passed to the **align** step. A small `--max-seqs` value can dramatically accelerate the entire alignment process, but may impair sensitivity when there are many hits; furthermore, there is an unintuitive relationship between prefilter score and final `nail` scores, meaning there is no straightforward way to select either `--max-seqs` or prefilter score thresholds that will preserve sensitivity while maintaining good speed.

While `nail` can be run with a fixed value for MMseqs2’s `--max-seqs` parameter, by default it runs **prefilter** without the standard `--max-seqs` per-query hard limit. Rather than passing the entire set of query-target pairs to **align**, `nail` uses an iterative scheme that adapts to each query’s hit density: queries are retired as soon as their productive matches run out, but allowed to continue indefinitely while productive matches are still being produced. The procedure is:

1. For each query, align the next *p* (default: 200) highest scoring pairs
2. For each query, compute the fraction *x* of those *p* pairs that pass `nail`’s seed filter and are sent to cloud search
3. Queries with *x > y* (default: *y* = 0.01) continue to the next batch; the rest are retired from seeding
4. Increase *p* (default: *p* ← 2*p*)
5. Repeat from step 1 until all queries are retired or pairs are exhausted

Queries with only a handful of homologs see *x* fall below *y* after one or two batches and are retired quickly. Queries with thousands of homologs maintain a high *x* across many batches, allowing `nail` to keep drawing from the prefilter’s score-sorted output until the productive matches taper off. A fixed `--max-seqs`, as is employed in MMseqs2, does not adapt this way: setting it aggressively risks cutting off queries that still have distant homologs available in the prefilter output, while a loose threshold can lead to excessive computation. This scheme stops only when the data themselves signal that the output has run out of productive candidates. See Supplementary Figure S1 for an analysis of progressive seeding and fixed max-seqs settings.

### Converting pHMMs to MMseqs2 profiles

Both MMseqs2 and HMMER are designed to search using a profile that captures position-specific scores based on a provided multiple sequence alignment (MSA) that represents a collection of related query sequences. The tools differ in the precise conversion of MSA-observed per-column residue counts into per-position scores. A greater challenge for `nail` is that the tools use different rules to decide which columns in the family MSA correspond to scoring positions in the learned profile (the tools generally select columns in which the majority of observed sequences contain a residue, but sequence weighting in HMMER differs from the approach in MMseqs2). To avoid the complexity of registering positions across these two rule-sets, `nail` directly creates an MMseqs2 profile by converting pHMM match state values to the MMseqs2 profile format.

### Test Environment

All tests were performed using 8 threads on a Linux workstation with an Intel i9-14900KF (6.0GHz boost) 24 core processor and 128GB RAM. Standard wall clock times were captured.

## Discussion

As implemented, `nail` demonstrates that it is possible to employ powerful Forward/Backward inference with significantly reduced time and memory requirements. Here, we highlight ways in which we expect future advances may lead to superior annotation performance.

### Better candidate seeds

`nail`’s dependency on MMseqs2 creates two common ways that a good alignment can be missed. In the most straightforward one, the MMseqs2 portion of the pipeline fails to find a good alignment candidate, so `nail`’s sparse Forward/Backward stage is never given a chance to identify the match. The fast k-mer match stage of MMseqs2 is the common cause of such misses, and is responsible for most of the sensitivity difference between `nail` and HMMER3. `nail`’s modular implementation makes it possible to explore development of new candidate detection options with no exposure to other parts of the algorithm. Fast and highly sensitive candidate detection may be improved through an alternative k-mer matching scheme (perhaps leveraging fast FM-index implementation as with 2), neural networks [34, 28], minimizer analogs [33, 22], hardware accelerators [3], or other methods.

### Reporting fragments or multiple domains

A more subtle issue is that the current `nail` pipeline only analyzes the MMseqs2-sourced region with the highest score; it does not explore lower-scoring MMseqs2 matches to identify a superior Forward/Backward score/alignment. The most common impact of this will be that only a single match will be reported when there are in fact multiple hits to be found, as will be true when there are multiple copies of a query domain, or a highly fragmented sequence match. Mechanisms for identifying multiple good begin/end seeds, and for efficiently managing the associated sparse cloud(s), will improve `nail`’s completeness and sensitivity.

The *colinear match* class characterized in Figure 6 is the principal target of this extension, and would close the part of the multi-domain recovery gap that corresponds to finding single instances of query family models.

### DNA search

`nail` will be extended to support DNA-to-DNA search, as in HMMER3’s `nhmmer` [40]. `nail` will also be extended to support annotation of protein-coding DNA, including models for both splicing activity and error-inducing frameshifts, as in our BATH software [23].

### Support for more complex models

`nail` reduces the computation workspace while retaining the core models of pHMM search. With this architecture in place, it will be possible to expand model complexity while retaining desirable run time properties. For example, once the search space has been reduced to a small cloud, it is possible to incorporate per-target-position scoring features into the algorithm, perhaps enabling direct incorporation of models of sequence repetition [14, 27] into alignment scoring.

### Faster computations

The Forward/Backward recurrence calculations are modeled after the generic implementation in HMMER, with significant overhead required to support movement back and forth to log-scaled representations of likelihood ratios. Dynamic scaling in probability space is faster [11] and is expected to roughly double the speed of the Cloud and Forward implementations.

## Author contributions statement

TJW and JWR conceived algorithms design. JWR implemented software based on a complete prototype implementation by DHR. DHR conducted pilot experiments; JWR and TJW collectively conducted all experiments presented in the manuscript. All authors analyzed results and wrote aspects of the manuscript; all authors reviewed, edited, and approved the final manuscript.

## Acknowledgments

We are grateful to Genevieve Krause for helpful discussions during development of software and benchmarks. We thank Clément Goubert for assistance with visualization of multi-domain classifications. We also gratefully acknowledge the computational resources and expert administration provided by the University of Montana’s Griz Shared Computing Cluster (GSCC), and the high performance computing (HPC) resources supported by the University of Arizona TRIF, UITS, and Research, Innovation, and Impact (RII) and maintained by the UofA Research Technologies department.

## Supplementary Materials

Additional supplementary materials can be found at https://github.com/TravisWheelerLab/nail-benchmarks, including code for construction and analysis of benchmarks.

### Max-seqs and Progressive seeding

Time versus recall for tool variants when searching Pfam against a single block of ~2.4 million MGnify sequences, exploring the importance of both the filter sensitivity setting (`-s`) and the maximum number of sequences passed through the MMseqs2 filter for full alignment (`--max-seqs`).

**Fig. S1:**
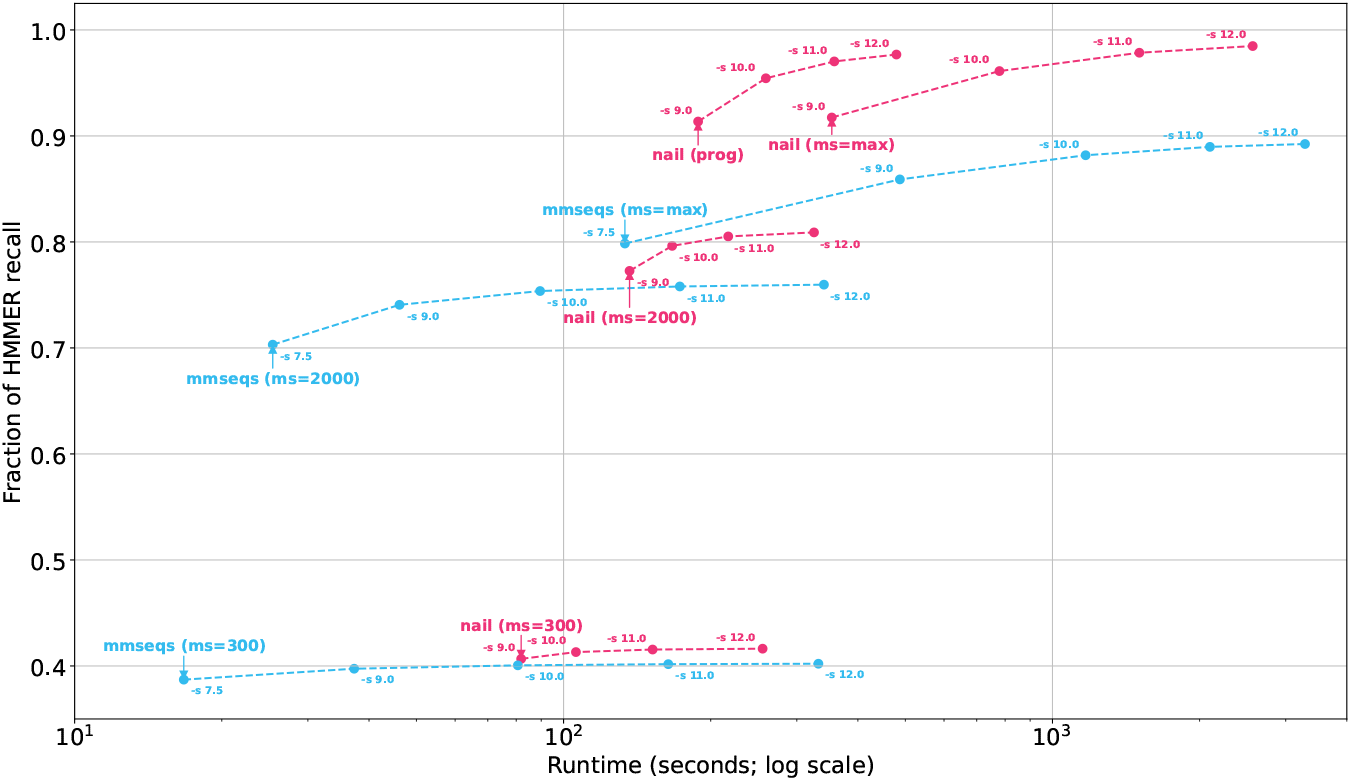
MMseqs2 was tested with three `--max-seqs` settings: 300 (default), 2000, and UINT MAX. For each of these max-seqs settings, it was run with five `-s` settings: 7.5 (default), along with 9.0, 10.0, 11.0, and 12.0); all points sharing the same max-seqs value are connected by lines. These lines show the recall gains and time penalty achieved by increasing both sensitivity and max-seqs settings. Similarly, `nail` variants was tested with `--mmseqs-s` at various settings (9.0, 10.0, 11.0, 12.0) and a sequence cap defined by either the default (progressive seeding) or overridden with `--mmseqs-max-seqs` set to either 300, 2000, or UINT MAX. The results demonstrate (1) that /nail offers improved recall over MMseq2 (consistent with Figure 4), (2) the importance of allowing a sufficient number of maximum sequences, and (3) the fact that `nail`’s progressive strategy achieves essentially identical results to a more complete run of `nail` with unbounded max, but with reduced runtime. Note: for any given input, it is not possible to guess the optimal choice for max-seqs without a prior expectation on the number of matches that will be found. All tools were run with 48 threads.

### Large-scale MGnify-Pfam annotation, additional analyses

All analyses in this section use the best-family-filtered MGnify-Pfam annotation defined in the main text.

`nail` performance for HMMER hits.

Table S1 reports, for each *n*_*pos*_ bin (the number of positive-scoring domain alignments HMMER reports for the pair), how HMMER’s hits are recovered by `nail`, grouped by whether at least one individual domain alignment exceeds the gathering threshold *h*_*cut*_. The three `nail` outcomes are: *pass* (score ≥ *n*_*cut*_), *sub* (score reported but *< n*_*cut*_), and *none* (no `nail` score for the pair; MMseqs2 may or may not have a hit). Percentages give `nail`’s outcome rate relative to HMMER hits in each cell. `nail` recovers 91% to 95% of HMMER hits supported by at least one strong individual alignment, across all *n*_*pos*_. When no single domain alignment clears threshold, `nail`’s pass rate falls to 31% to 41%, reflecting that `nail` does not aggregate sub-threshold domain evidence.

### Category counts

Table S2 reports the underlying counts for Figure 6, from a grouped sample of HMMER-only multi-domain pairs capped at 50 (family, target) pairs per Pfam family.

**Table S1.** nail outcomes for all HMMER hits in the MGnify-Pfam annotation, grouped by *n*_*pos*_ and by whether any individual domain alignment exceeds the gathering threshold *h*_*cut*_. Percentages give the outcome rate relative to HMMER hits in each column; HMMER rows are the column denominators. Counts are rounded to the nearest thousand.

| $n_{pos}$ | outcome | $\geq 1 \text{ dom} > h_{cut}$ | | no dom $> h_{cut}$ | | total | |
| --- | --- | --- | --- | --- | --- | --- | --- |
|  |  | count | % | count | % | count | % |
| 1 | HMMER (all) | 1,877,206,000 |  | 62,059,000 |  | 1,939,265,000 |  |
|  | nail pass | 1,788,884,000 | 95.3% | 29,931,000 | 48.2% | 1,818,815,000 | 93.8% |
|  | nail sub | 9,260,000 | 0.5% | 18,163,000 | 29.3% | 27,423,000 | 1.4% |
|  | nail none | 79,062,000 | 4.2% | 13,965,000 | 22.5% | 93,027,000 | 4.8% |
| 2 | HMMER (all) | 143,567,000 |  | 96,141,000 |  | 239,708,000 |  |
|  | nail pass | 136,457,000 | 95.0% | 15,475,000 | 16.1% | 151,932,000 | 63.4% |
|  | nail sub | 2,168,000 | 1.5% | 52,759,000 | 54.9% | 54,927,000 | 22.9% |
|  | nail none | 4,942,000 | 3.4% | 27,907,000 | 29.0% | 32,849,000 | 13.7% |
| 3 | HMMER (all) | 18,326,000 |  | 26,033,000 |  | 44,359,000 |  |
|  | nail pass | 17,104,000 | 93.3% | 4,000,000 | 15.4% | 21,104,000 | 47.6% |
|  | nail sub | 354,000 | 1.9% | 12,104,000 | 46.5% | 12,458,000 | 28.1% |
|  | nail none | 868,000 | 4.7% | 9,929,000 | 38.1% | 10,797,000 | 24.3% |
| 4 | HMMER (all) | 7,187,000 |  | 12,846,000 |  | 20,033,000 |  |
|  | nail pass | 6,586,000 | 91.6% | 2,002,000 | 15.6% | 8,588,000 | 42.9% |
|  | nail sub | 129,000 | 1.8% | 5,886,000 | 45.8% | 6,015,000 | 30.0% |
|  | nail none | 472,000 | 6.6% | 4,958,000 | 38.6% | 5,430,000 | 27.1% |
| 5 | HMMER (all) | 3,385,000 |  | 6,077,000 |  | 9,462,000 |  |
|  | nail pass | 3,078,000 | 90.9% | 1,049,000 | 17.3% | 4,127,000 | 43.6% |
|  | nail sub | 39,000 | 1.1% | 2,696,000 | 44.4% | 2,735,000 | 28.9% |
|  | nail none | 269,000 | 7.9% | 2,332,000 | 38.4% | 2,601,000 | 27.5% |
| 6 | HMMER (all) | 2,262,000 |  | 3,224,000 |  | 5,486,000 |  |
|  | nail pass | 2,069,000 | 91.5% | 647,000 | 20.1% | 2,716,000 | 49.5% |
|  | nail sub | 24,000 | 1.0% | 1,399,000 | 43.4% | 1,423,000 | 25.9% |
|  | nail none | 169,000 | 7.5% | 1,178,000 | 36.5% | 1,347,000 | 24.6% |
| $\geq 7$ | HMMER (all) | 3,628,000 | | 4,518,000 | | 8,146,000 | |
|  | nail pass | 3,224,000 | 88.9% | 1,068,000 | 23.6% | 4,292,000 | 52.7% |
|  | nail sub | 38,000 | 1.0% | 1,779,000 | 39.4% | 1,817,000 | 22.3% |
|  | nail none | 366,000 | 10.1% | 1,671,000 | 37.0% | 2,037,000 | 25.0% |

**Table S2.** Counts of HMMER multi-domain hits that are missed by nail, grouped by *n*_*pos*_ and category. These numbers represent a sample taken from the full hit set, in which each Pfam family contributes at most 50 (family, target) pairs per Pfam family; 1,239,880 pairs total).

| category | $n_{pos} = 2$ | $n_{pos} = 3$ | $n_{pos} = 4$ | $n_{pos} = 5$ | $n_{pos} = 6$ | $n_{pos} \geq 7$ | total |
| --- | --- | --- | --- | --- | --- | --- | --- |
| *strong_match* | 39,454 | 9,424 | 3,731 | 2,512 | 2,017 | 2,893 | 60,031 |
| *colinear_match* | 162,293 | 35,424 | 4,747 | 785 | 234 | 84 | 203,567 |
| *pileup:full* | 61,700 | 32,539 | 23,348 | 16,449 | 13,933 | 13,793 | 161,762 |
| *pileup:mid* | 50,631 | 39,713 | 29,681 | 21,273 | 15,633 | 17,304 | 174,235 |
| *pileup:fragment* | 74,168 | 122,111 | 115,536 | 87,471 | 61,399 | 70,659 | 531,344 |
| *shuffled_match* | 28,729 | 6,882 | 805 | 190 | 57 | 18 | 36,681 |
| *weak_match* | 16,334 | 16,308 | 7,839 | 8,876 | 9,335 | 13,568 | 72,260 |
| total | 433,309 | 262,401 | 185,687 | 137,556 | 102,608 | 118,319 | 1,239,880 |

### Per-copy strength within each category

The classification above captures the spatial arrangement of domain hits but does not say how strong any single hit is. A *pileup:fragment* call where the best copy scored just below *h*_*cut*_ differs qualitatively from one whose strongest copy is far below threshold: in the first case, HMMER’s call is anchored by a near-threshold alignment with accumulation supplying the small remainder, while in the second, the call rests entirely on summing low-quality fragments. Table S3 reports, for each category and *n*_*pos*_ bin, the fraction of cases for which the maximum single-domain bit score (dom_max) comes within 10 bits of *h*_*cut*_. By construction, dom max *< h*_*cut*_ in every non-*strong match* category; values close to 100% indicate that most cases in that cell are anchored by a near-threshold copy, while values close to 0% indicate that the call rests on accumulation alone. The contrast is sharpest in *pileup:fragment* at *n*_*pos*_ ≥ 4: only 2% to 6% of these cases have any single domain alignment within 10 bits of threshold, so the call rests on summing many weak fragments rather than on a near-threshold copy plus accumulation.

**Table S3.**
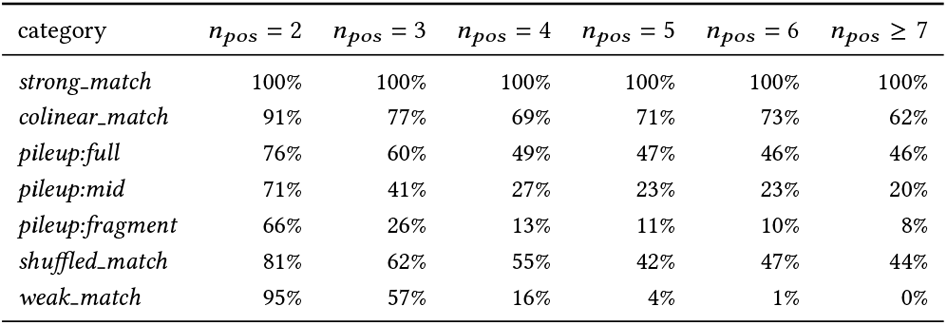
Percentage of cases with dom_max ≥ *h*_*cut*_ − 10, by category and *n*_*pos*_. Hyphens indicate categories with fewer than 10 sample cases in the bin.

| category | $n_{pos} = 2$ | $n_{pos} = 3$ | $n_{pos} = 4$ | $n_{pos} = 5$ | $n_{pos} = 6$ | $n_{pos} \geq 7$ |
| --- | --- | --- | --- | --- | --- | --- |
| *strong_match* | 100% | 100% | 100% | 100% | 100% | 100% |
| *colinear_match* | 91% | 77% | 69% | 71% | 73% | 62% |
| *pileup:full* | 76% | 60% | 49% | 47% | 46% | 46% |
| *pileup:mid* | 71% | 41% | 27% | 23% | 23% | 20% |
| *pileup:fragment* | 66% | 26% | 13% | 11% | 10% | 8% |
| *shuffled_match* | 81% | 62% | 55% | 42% | 47% | 44% |
| *weak_match* | 95% | 57% | 16% | 4% | 1% | 0% |

### Classification algorithm

For each HMMER-only multi-domain hit, the spatial arrangement of its domain alignments on the HMM is classified by the following procedure. Categories are tested in priority order; the first match is returned. We acknowledge that this is a classification heuristic, and note that there is no established method for classifying matches according to their complex sets of overlapping hit fragments; we note that this strategy anecdotally assigns classes that usually agree with our manually assigned labels.

1. If any individual domain bit score exceeds *h*_*cut*_: return *strong_match*.
2. Filter to *important* domains, defined as bit score ≥ 4.0. Sub-4-bit hits contribute negligibly to sequence score and add noise to spatial analysis. If no important domains remain: return *weak_match*.
3. Build pileups of mutually-overlapping important domains. Two hits are connected if their HMM-space intersection exceeds 50% of the shorter span; pileups are connected components found by union-find. An iterative splitting step handles cases where one domain is much stronger than the pileup core: the strong outlier is split at the pileup’s majority HMM range, with the core slice absorbed into the pileup and the flanking overhangs returned to the unassigned pool if they are at least 20 residues long. Termination is guaranteed because each split strictly reduces the total HMM span of unassigned domains.
4. Compute (a) the maximum-weight co-linear chain score by dynamic programming, where two hits may chain if they are ordered consistently in both HMM and target space (small boundary overlaps of up to 10 residues and up to 10% of the shorter span are tolerated), and (b) the unordered sum of per-pileup maximum scores.
5. If the best multi-element pileup’s score exceeds the chain score, return *pileup:full, pileup:mid*, or *pileup:fragment* according to whether the pileup’s majority HMM range covers ≥70%, 50% to 70%, or *<*50% of the HMM.
6. Otherwise, if the chain score reaches *h*_*cut*_ − 4: return *colinear match*.
7. Otherwise, if the unordered score reaches *h*_*cut*_ − 4: return *shuffled_match*.
8. Otherwise: return *weak_match*.

The 4-bit tolerance in steps 6 and 7 accounts for the importance filter in step 2: without it, many borderline cases would fall through to *weak_match* only because small contributions were excluded.

